# Co-occurring Mutations in Different Genes Can Fuel Oncogenic Signaling and Serve as Metastatic Tumor Markers

**DOI:** 10.1101/2024.05.01.592039

**Authors:** Bengi Ruken Yavuz, Ugur Sahin, Hyunbum Jang, Ruth Nussinov, Nurcan Tuncbag

**Affiliations:** Cancer Innovation Laboratory, National Cancer Institute, Frederick, MD, 21702, USA; Department of Molecular Biology and Genetics, Koç University, Istanbul, Turkey; Computational Structural Biology Section, Frederick National Laboratory for Cancer Research, National Cancer Institute at Frederick, Frederick, MD 21702, USA; Department of Human Molecular Genetics and Biochemistry, Sackler School of Medicine, Tel Aviv University, Tel Aviv 69978, Israel; Department of Chemical and Biological Engineering, College of Engineering, Koç University, Istanbul, Turkey; School of Medicine, Koç University, Istanbul, Turkey; Koc University Research Center for Translational Medicine (KUTTAM), Istanbul, 34450 Turkey

**Keywords:** oncogene induced senescence (OIS), mutation doublets, molecular signatures of cancer, cancer genome analysis, cancer type specific subnetworks, metastatic markers

## Abstract

Interrogation of big genomic data and integration with large-scale protein-protein interaction networks and pathways, can provide deep patterns that are rare– yet can prompt dramatic phenotypic alterations and serve as clinical signatures. Mapping cancer-specific co-occurring mutation-pair signatures, in primary and metastatic tumors, is indispensable in precision oncology. The additivity of co-occurring driver mutations in different genes (*in trans*) can lead to powerful proliferation signals. Co-occurring rare *in trans* combinations can serve as metastasis markers; excluded combinations may indicate candidates for oncogene-induced senescence (OIS), a tumor-suppressive mechanism. Our statistical framework of the pan-cancer mutation profiles of ∼60,000 tumor sequences from the TCGA and AACR GENIE databases, identified 3424 statistically significant different double mutations in non-redundant pathways, that is, have different downstream targets that may promote specific cancers through single or multiple pathways. Our analysis indicates that they are mostly in primary tumors. We list actionable *in trans* mutations for 2385 metastatic tumors and provide co-occurrence trees of metastatic breast-cancer markers. This innovative work clarifies the mechanistic conceptual basis and establishes the first of its kind tool for identifying and predicting metastasis. Crucially, when coupled with their proliferative functions and pathways, and linked with drugs, it could provide an invaluable metastasis-targeting resource.

## Introduction

Tumorigenesis is a complex process, which could be attributed to alterations in a group of genes(1,2). Co-occurrence, or conversely, exclusivity, of two mutations in different genes (i.e., *in trans*) could be a random event in cancer evolution, which is then selected or excluded to provide a functional advantage. If two mutations in different genes occur less or more frequently than expected, they are mutually exclusive or co-occurring, respectively (3,4). Mutual exclusivity was mostly observed in genes in the same or redundant pathways, and conversely, a co-occurrence trend was observed in genes in different, or parallel pathways (5). In our definition, if the pathways recruit the same downstream protein families, they are redundant; if evolutionary-independent, they are parallel. Exclusivity has been attributed to the expectation that acquiring a second strong mutation in the same pathway is unsustainable for the cell since it risks its death (3,4). Excluded co-occurrence has been proposed as associated with tumor subtype, synthetic lethality, and positive selection(6). We clarified that co-occurrence is restricted (7,8), since a sustained, additive effect on signaling strength of same — or redundant— pathway driver mutations is likely to hyperactivate the proliferation signal, triggering an oncogene-induced senescence (OIS) cellular program (9–11). Phenomenologically, ‘synthetic lethality’, where a strong mutation in one gene product can be sustained but not in two, resembles such excluded OIS co-occurrence. Signaling strength can provide its molecular basis.

OIS is a mechanism the cancer cell uses to proliferate- yet dodge cell death. Cancer cells select combinations of potent mutations. As early as 1953 (12) , and again in 1969 (13), 1971 (14), 1999 (15), 2002 (16), 2015 (17), and more recently, it has been established that multiple co- existing mutations are required for the emergence of cancer. We also know that the number increases during proliferation. Yet, cells survive, suggesting that OIS combinations were sidestepped, or more likely, since somatic mutations emerge sporadically, cells harboring excluded combinations did not endure (18). Activating mutations in genes encoding RAS- and RAF-family proteins, especially the strong *BRAF^V600E^* hotspot is one example (19,20). Co- expression of *KRAS*^G12D^ and *BRAF*^V600E^, two strong drivers, led to OIS in lung cancer in mouse models (21). The absence of co-occurring *NRAS* and *BRAF* variants, such as *BRAF^V600E^*, in the RTK-RAS pathway in the TCGA skin cutaneous melanoma cohort illustrates mutual exclusivity. Similarly, *BRAF* and RAS gene family mutations are mutually exclusive in metastatic colorectal cancer tumors (19). OIS, a tumor-suppressive mechanism arresting cell cycle progression, can be the main reason for excluding the co-occurrence of driver mutations in the same, here MAPK, pathway (22–24). Irreversible senescence phenotypes express strong oncogenic stimuli (25). Along similar lines, overexpression of oncogenes such as *KRAS*, *BRAF*, and *MYC*, which also generate strong proliferation signal, induce OIS, as does increased PI3K/AKT signaling. Loss of *PTEN*, which negatively regulates the PI3K/AKT pathway, can trigger OIS through a p53- dependent pathway (26), or via coupling with strong PI3K mutational variants. Both constitutively generate signaling lipid PIP_3_. Sustained hyperactivation of the PI3K/AKT/mTORC1 pathway results in OIS (27,28). Examples of mutually exclusive relations between gene pairs include *BRCA2*-*TP53*, *BRCA1*-*PARP1*, and *PTEN*-*PIK3CA* in breast cancer; in ovary cancer *BRCA1*-*CCNE1*, *BRAF*-*KRAS*, *ERBB2*-*KRAS* (*29*). A functional survey of oncogene-tumor suppressor linkage in mouse models of lung tumors revealed that tumor suppressor inactivation often alternates between advantageous and detrimental contingent upon the oncogenic context (30). In all cases, overexpression and strong mutational variants behave similarly since both result in a large number of protein molecules in their active, catalysis-primed state. Comprehensive modeling of mutations and copy number alterations (CNAs) offered that a third mutation, with the hit in the same— or different— gene in the same pathway can reflect alternate evolutionary pathways to cancer (31), though likely still harnessing this signaling premise.

In addition to prediction of driver mutations (32,33), identifying viable evolutionary trajectories also requires systematic investigation of common pairwise and higher-order epistasis between mutations in cancer genomes (6,32,34–36). There are several tools for identifying biologically meaningful, mutually exclusive gene sets (29), including statistical frameworks (MEScan (37), MEGSA(38), CoMET(39)) and driver module detection (MEMo (38,40), CoMDP (41)). SELECT captures evolutionarily dependent alterations that can affect drug resistance and sensitivity at the pan-cancer level (42).

Complex biological functions are mostly managed through PPIs, and mutations occurring in proteins may alter the cell phenotypes (43). Alterations perturbing PPIs can impact drug outcomes(44–46). Detection of druggable oncoPPIs(44) is promising since some could constitute more cancer-specific therapeutic targets (47,48), although the pre-existing mutation load can impact the efficacy (49). Among the Fujian cohort mutations classified as metastatic markers in gastric carcinomas, one is a *PTPRT* mutation (50), and another, Chondrosarcoma *TERT* promoter mutation is a metastasis marker(51). A real-world clinicogenomic dataset permitted the discovery of biomarkers that predict treatment outcomes that affect patients’ survival (52). Treatment-specific genetic alterations in non-small cell lung cancer (NSCLC) include mutually exclusive and co-occurring mutations (53,54).

In this study, we identified 3424 different gene double mutations on ∼60,000 tumors. Doublets *in trans* mutations are usually composed of a frequent and a rare mutation. Mutations disrupting similar functions are mutually exclusive and genes harboring different double mutations are mostly populated in different pathways. As to metastatic tumors, some double mutations *in trans* were significantly higher, and these could serve as metastatic markers, and help address the potential mechanisms which are involved. The mutational burden of primary tumors can be significant and grow over time. Surprisingly, this does not appear to be the case for metastatic tumors, whose mutational burden increase is more moderate (55), raising the question of *how metastatic phenotypes arise and whether their genomes hold clue*s. We believe that it likely involves emerging OIS. The availability of metastatic samples prompted us to investigate co-occurrence patterns of mutations enriched in metastatic cancers and compare them with those of primary breast cancer. Precision medicine efforts can benefit from understanding the relationships between epistatic interactions and treatment response. We provide a list of actionable mutations for metastatic tumors in breast cancer cohort and a tree of double, triple, and quadruple co-occurrence patterns across the pan-cancer metastatic tumors. They also address the questions of why these signatures, and what can they tell us about the possible the molecular mechanisms which are involved. However, more complete data and analysis are needed.

## Materials and Methods

### Data collection and processing

All available somatic missense mutation profiles are downloaded from The Cancer Genome Atlas (TCGA) and the AACR launched Project GENIE (Genomics Evidence Neoplasia Information Exchange) (56,57). The TCGA mutation annotation file contains more than 10,000 tumors across 33 different cancer types. We used the merged MC3 file to get TCGA pan-cancer data. The somatic variants without sufficient depth coverage and variants found in the panel of normal samples were evaluated as possible germline variants and were removed from the file before merging.

The GENIE mutation file (Release 6.2-public) contains 65,401 patients and 68,897 tumor samples across 648 cancer subtypes under the Oncotree classification. Within the GENIE cohort 2930 patients match with multiple tumor barcodes. For those cases, only one primary tumor barcode is kept when available; if not, only one metastatic or unspecified tumor barcode is kept for further analysis without any other constraint. Among these patients, 2019 has sequenced primary tumors, 757 have sequenced metastatic tumors and the remaining 154 have tumors of the type not specified.

Next, we selected non-synonymous mutations including missense, nonsense, nonstop and frameshift mutations (altering only one position on a protein). We also excluded mutations where the wild type and/or mutant residue name is not specified. As a result of this filtering process, 9703 and 57,921 tumors remained with a total of 1,631,755 point mutations in the TCGA and GENIE cohorts, respectively.

We first pre-filtered on the VAF (Variant Allele Frequency) value to control the heterogeneity of the samples to some extent given that variants were collected by bulk sequencing in both datasets. We calculated VAF by using the ratio of the values in the t_alt_count and t_depth columns of the MAF (Mutation Annotation File) file of the pan-cancer data sets. We continued our analysis with the mutations that have a VAF value more than 0.125, ensuring that the remaining mutations are present roughly in 25% of the sequenced cells. We identified 3519 tumors as hyper-mutated out of 62,567 samples with at least one point-mutation with a Q3+8 x IQR threshold.

We continued the analyses with 59,048 non-hyper-mutated samples from 619 cancer subtypes and 33 tissues (including OTHER category).

### Identification of Significant Double Alterations

The total number of mutations with VAF>0.125 is 395,801; among these mutations 20,157 are observed in three or more non-hyper-mutated tumor samples. We constructed binary combinations of these 20,157 mutations. We created a contingency table for each possible binary combinations of 20,157 mutations (except for the mutations in the same gene) composed of tumor numbers having both mutations, only the first or second alteration and none of those two alterations among the 59,048 non-hyper-mutated samples. As a result, we obtained 366,573 potential different-gene double mutations to be tested in 59,048 non-hyper-mutated tumor samples and assessed their statistical significance (Fisher’s Exact Test). For each potential double mutation, we created a contingency table [[a,b],[c,d]] where a is the number of tumors having both alterations, b is the number of tumors having only the first alteration, c is the number of tumors having only the second alteration and d is the number of all tumors not having these two alterations together (d = 59,048-(a+b+c)). Here, we did not apply any pre-filtering on the number of double mutant tumors before conducting the Fisher’s Exact Test, i.e., the cases where a=0 are also tested for significance. Then, we applied multiple corrections by using Benjamini-Hochberg method and continued subsequent analyses with 3424 doublets having q<0.3 and observed in at least three tumors. We used the Catalog of Validated Oncogenic Mutations from the Cancer Genome Interpreter(58) to label double mutation components: if a mutation is among the 5601 driver mutations, we label it as known driver (D) if it is cataloged on the Cancer Genome Interpreter, otherwise passenger (P).

### Pathway Analysis

We used 46 signaling pathways from KEGG: Kyoto Encyclopedia of Genes and Genomes(59) to associate with double mutant genes. We downloaded the relevant data from https://maayanlab.cloud/Enrichr/.

### Oncoprint Maps

To reveal mutual exclusivity and co-occurrence patterns between double mutations we plotted oncoprint maps by using ComplexHeatmap(60) package of R and cBioPortal (https://www.cbioportal.org).

### Transcriptome Analysis

To identify differentially expressed genes in the group of patients with double mutations compared to the single mutant patient group, we downloaded RNA-seq transcriptome data of the TCGA project from the cBioPortal database (https://www.cbioportal.org). We used median Transcripts Per Kilobase Million (TPM) values of RNA-seq data of PAAD cohort of TCGA. For the PAAD cohort, 177 patients with TPM values, we constructed two groups, where Group 1 is tumors having at least one significant double mutation of type KRAS^G12D/V/C^ +TP53^mutation^ and Group 2 is tumors having either single mutant KRAS^G12D/V/C^ or single mutant TP53. We calculated the log_2_FC value of each gene between the double mutant and single mutant groups by using the formula:

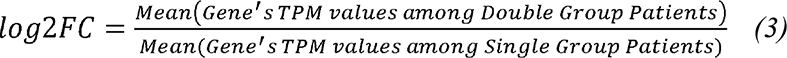

We identified differentially expressed genes between the double mutant tumors group and the single mutant tumors group (comparison of means of TPM values by Mann-Whitney U-Test). If |log2F[|>0.5 and p-value < 0.01 we considered the corresponding gene as differentially upregulated or downregulated in the double mutant group. We continued our analysis with these differentially expressed genes, and calculated z-scores of each gene as follows:

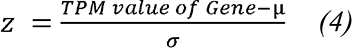

where is µ is the mean of TPM values and σ is the standard deviation across all samples in the double mutant and the single mutant groups. After obtaining z-scores for each gene, we sorted the genes as downregulated and upregulated and represented these values as a heatmap (https://seaborn.pydata.org).

We used Webgestalt (http://www.webgestalt.org) for the gene set enrichment analysis where the functional database is selected from Reactome and significantly up- or down-regulated pathways are found. FDR threshold is selected 0.05 and the list of genes ranked by their logFC values are given as input.

### FPGrowth Tree Construction

We are inspired by the prediction of association rules in database transaction systems which find the most frequent associations between items in each transaction. In our setup, each tumor is considered as a transaction and each alteration is considered as an item in the transaction. We used the mlextend library’s FreqItems and AssociationRules functions for mining frequent item sets and association rules. The FP growth algorithm is selected for tree construction where each node represents one alteration and each edge in the tree represents the association of the nodes (61). In the constructed tree, all nodes in the path from root to the distant node are associated with each other and strongly present together in the tumors. The implementation of this workflow or additional documentation on how the FP-Tree algorithm is adapted for this specific use case is available at https://github.com/ugur0sahin/FMPSeeker.

The tendency of the alterations to be specific to metastatic tumor is calculated by

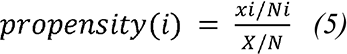

where *x_i_*is the number of metastatic tumors having double mutation *i*, *N_i_*is the number of tumors having double mutation *i*, X is the number of metastatic tumors and N is the number tumors in our dataset.

### Personalized PageRank Algorithm

We used a network diffusion-based algorithm to find the most affected region of the interactome given a set of nodes. Given a directed or undirected graph G(v,e) where v∈V and e∈E and a set of seed nodes S⊆V, the personalized PageRank algorithm solves the seed set expansion problem, where it finds which additional nodes may exist in the community besides the nodes in S and ranks them according to their importance. We used the PageRank (62) function implemented in Python networkx library (63). The damping parameter, alpha, is selected as 0.85 and the number of iterations are 100.

### Tissue Specific Subnetworks

To determine tissue specific seed genes to perform Personalized PageRank algorithm for obtaining tissue specific subnetworks, we calculated the fraction of double-mutant tumors among all tumors in the corresponding tissue for each doublet. Then we evaluated the double mutated gene couples’ components as seed genes if the fraction of the doublet is greater than 25%. We assigned initial weights as 1. We run the PageRank algorithm on the PPI network from OmniPath(64) with alpha = 0.85. We added the edges with score greater than the threshold 0.001 to the breast specific subnetworks.

We evaluated 46 signaling pathways from KEGG. We obtained pathway-double-mutant tumors by taking the union of all double mutants when at least one of the genes belong to the corresponding pathway. We filtered the pathways that have less than 80 double mutant tumors and tissues and pathways that have less than 10% double mutant tumor rate. We constructed cancer type specific subnetworks for breast and pancreas tissues with seed genes that have double mutations and tissue specific double mutant fraction is greater than a certain threshold (thresholds are as follows: breast, 0.5; pancreas, 1.5). The number of seed genes for breast cancer and pancreas cancer are 12 and 23, respectively.

## Results

### Co-occurrence of frequent and rare mutations in different genes in tumorigenesis

The tumor evolutionary selection of survival mutations and the intricate relationship between the initial mutations and tumor initiation, and the additional accumulating mutations and the more complex metastasis scenario, is still not fully understood, especially not the metastasis (65). Several studies tackled the initiation of the metastatic cascade at the clinical or molecular levels. Genomic features associated with metastasis have been identified for specific target organs as well as the genomic level differences between primary and metastatic tumors (50,66,67). It is crucial to identify distinctive and common genetic features of primary and metastatic tumors for personalized treatment strategies to lower the chance of drug resistance. To determine the contribution of the metastatic samples to the overall samples in the corresponding tissues, we first evaluated the number of metastatic tumors in the prefiltered pan-cancer data (see Methods).

The weight of metastatic tumors in tissues with at least 50 samples is shown in Figure 1A. Skin, breast, intestine, lung, ovary, and prostate are among the tissues that have relatively more metastatic cancers, despite the total number of metastatic tumors being less than primary tumors. To analyze the relation between mutations and cancer phenotypes more systematically, we identified statistically significant double mutations in different genes in ∼60,000 tumor samples from TCGA and GENIE Pan-Cancer datasets.

**Figure 1.**
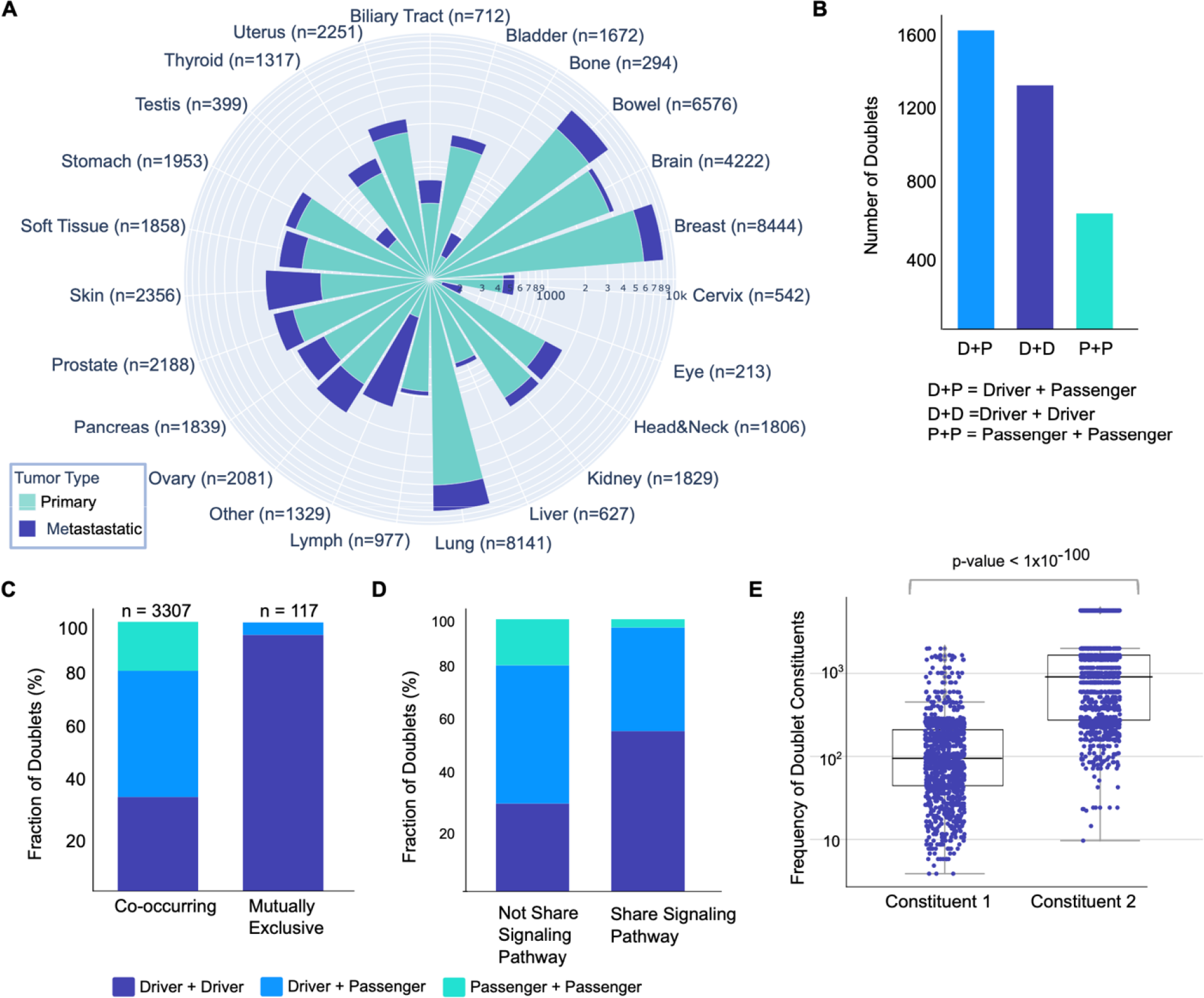
Overview of different gene double mutations. (A) The windrose plot shows the number of primary (blue) and metastatic (light blue) tumors among tissues with at least 50 samples. The number of all tumors for each cancer type is specified in parenthesis. Breast, bowel, and lung have the highest number of primary tumors. Although the total number of metastatic tumors are less than primary tumors overall, skin, breast, intestine, lung, ovary, and prostate are among the tissues that have relatively more metastatic cancers. **(B)** There are 3424 significant different gene double mutations (Fisher’s Exact Test p<0.05, Benjamini-Hochberg q<0.3). The doublet constituents are D+P (n=1554), D+D (n=1268) and P+P (n=602) where D is “Driver” and P is “Passenger” mutation. **(C)** Stacked bar plot of the different gene double mutation constituents for 3307 co-occurring and 117 mutually exclusive doublets. For co-occurring doublets, the distribution is as follows: D+D = 1155, D+P = 1550, P+P = 602. For mutually exclusive doublets there are 113 doublets of type D+D and 4 D+P doublets. D (Driver), P (Passenger). **(D)** We call a gene pair A|B if there is a different gene double mutation where the component mutations are on the genes A and B. We divided the different gene double mutations into two groups where gene pairs form these doublets: if the pair belong to any common pathway, we put the corresponding doublet into “Shared signaling pathway”, otherwise “Not shared signaling pathway”. 1280 doublet constituent genes share at least one signaling pathway where the corresponding doublets are of type: 750 D+D, 487 D+P, 39 P+P). 638 of them do not share a signaling pathway, and 206 doublets are of type D+D, 324 D+P, 108 P+P. For the remaining doublets, at least one of the genes harboring them does not have pathway information. Co-occurring different gene doublets are usually present on the gene pairs that are residing in different signaling pathways and this could imply some driver mutations need a helper passenger mutation from a different signaling pathway to increase oncogenic signaling. **(E)** Box plot showing the frequency of double mutation constituents. For each doublet, the component with lower frequency is put into “Constituent 1” and the other into “Constituent 2”. Regardless of being a driver or passenger mutation, most doublets are formed by combination of a frequent and a rare mutation. The figure shows that most of the doublets are composed of a frequent and a rare mutation.

In our analyses, we considered all non-synonymous mutations, including frameshift, nonsense, missense, and nonstop mutations. Frameshifts (insertions or deletions) that change a protein at more than one location were disregarded. Variants in which the wild type and/or mutant residues are not defined were also eliminated. We further eliminated mutations with VAF (Variant Allele Frequency) of less than or equal to 0.125 to ensure that the mutations are present in roughly 25% of the tumor cells that were sequenced. This yielded 62,567 samples and 19,415 genes carrying point mutations. Then we removed Hypermutated samples during preprocessing; there are 59,048 and 3519 non-hypermutated and hypermutated samples, respectively. We determined 366,573 potential different gene double mutations by forming binary combinations of the 20,157 point mutations that are mutated in at least three non-hypermutated tumors.

Collecting missense mutations in each gene and counting their pairwise combinations resulted in 3424 significant double mutations in different genes composed of co-existing mutations from 285 genes (Fisher Exact Test p<0.05, Benjamini-Hochberg q<0.3) (Supplementary Table 1). We annotated the significant doublet components, driver or passenger mutations. We used the Catalog of Validated Oncogenic Mutations from the Cancer Genome Interpreter (58) to label different gene double mutation components: if a mutation is among the 5601 driver mutations, we labeled it as a driver (D), otherwise passenger mutation (P). Among 3424 significant doublets, the number of doublets with constituents of type D+P, D+D, and P+P are 1554, 1268, and 602, respectively (Figure 1B). We also evaluated the co-occurrence and mutual exclusivity of doublets components and observed that 3307 of the doublets are co-occurring and 117 are mutually exclusive (Figure 1C, Methods). Among the 3307 co-occurring doublets, the number of D+D, D+P, and P+P pairs are 115, 1550, and 602, respectively. Similarly, for 117 mutually exclusive doublets 113 are of type D+D and four are of type D+P.

We represent a gene pair as A|B if there is a different gene double mutation where the component mutations are in genes A and B, and the gene pair A|B is mutated among the union of tumors with double mutations in these genes. 1280 gene pairs are carrying at least one significant doublet. We checked whether the genes comprising the gene pairs belong to the same signaling pathway. To carry out this analysis we retrieved 46 signaling pathways and harbored at least one different gene double mutation from KEGG. In most cases, a gene belongs to more than one signaling pathway. We evaluated gene pair components in terms of sharing (or not sharing) at least one pathway and observed that 1280 gene pair constituents share at least one signaling pathway (750 D+D, 487 D+P, 39 P+P), and 638 of them (206 D+D, 324 D+P, 108 P+P) do not share (Figure 1D). Moreover, we compared the distributions of doublet constituents of each different gene double mutation (Figure 1E, Figure S1). For each doublet, we put the constituent with lower frequency into the “Constituent 1” category and the other into the “Constituent 2” category. Comparing the frequencies of the mutations in each category we see that most of the doublets are composed of a frequent and a rare mutation.

Among the different double mutations, odds ratio analysis shows that most of the doublets are co-occurring (n=3307) and within this group, ∼45% are composed of a driver and a passenger mutation. These types of doublets also constitute a large portion among the doublets that do not share a signaling pathway. This shows that co-occurring different gene doublets are usually harbored by the gene pairs that are residing in different signaling pathways and this could imply some driver mutations need a helper passenger mutation from a different signaling pathway to increase oncogenic signaling. On the other hand, almost all mutually exclusive doublets are combinations of driver mutations. For each double mutation, the constituents are usually frequent and rare implying regardless of the mutation type (being passenger or driver) an intense oncogenic signaling is facilitated via the help of a less frequent mutation.

### Some gene pairs co-mutate in certain tissues

Mutations exist in normal tissues too (68). Cancer can develop as mutations accumulate in tissues over time where their contribution to the tumor progression varies across different classes of mutations (69). These mutations give cancer cells unique properties referred to as the cancer hallmarks. Most cancer driver genes are altered in certain cancers but not in others, indicating their sporadic (rather than germline) nature, thus tendency to be tissue, or cell type, dependent. Consistently, due to chromatin organization, different cell types differ in their expressed proteomes, the chance of a therapeutic success is frequently influenced by the type of malignancy (70,71). Dedifferentiation within the cell’s lineage can transition the original cancer to others, as shown in adagrasib resistance (49). In a previous study, we identified latent driver mutations through double mutations in the same gene (in *cis*) that are specific to certain cancers (72). Here we discover double mutations in different genes (*in trans*) and the tissues and cancer subtypes they are most prominently populated in.

Our statistical framework yielded 1284 gene pairs composing 3424 different gene doublets. We demonstrate the tissue enrichment of the gene pairs where both components are oncogenes (Figure 2A) and when one is oncogene (OG) while the other one is a tumor suppressor gene (TSG) (Figure 2B). Node sizes represent the number of tumors having mutations in gene pair mutant tumors in the corresponding tissue. We obtained the gene pair mutants by aggregating all tumors carrying different gene double mutations in the corresponding gene pair components. The size of the samples with mutated gene A or B in each tissue was used as a denominator to calculate the fraction represented in the node colors. To simplify Figure 2A, only oncogene pairs and tissues with more than 10 double mutations are kept (47 gene pairs). Similarly for Figure 2B, OG and TSG pairs with more than 20 double mutations are kept (54 gene pairs). Same filtering applied to the tissues as well.

**Figure 2.**
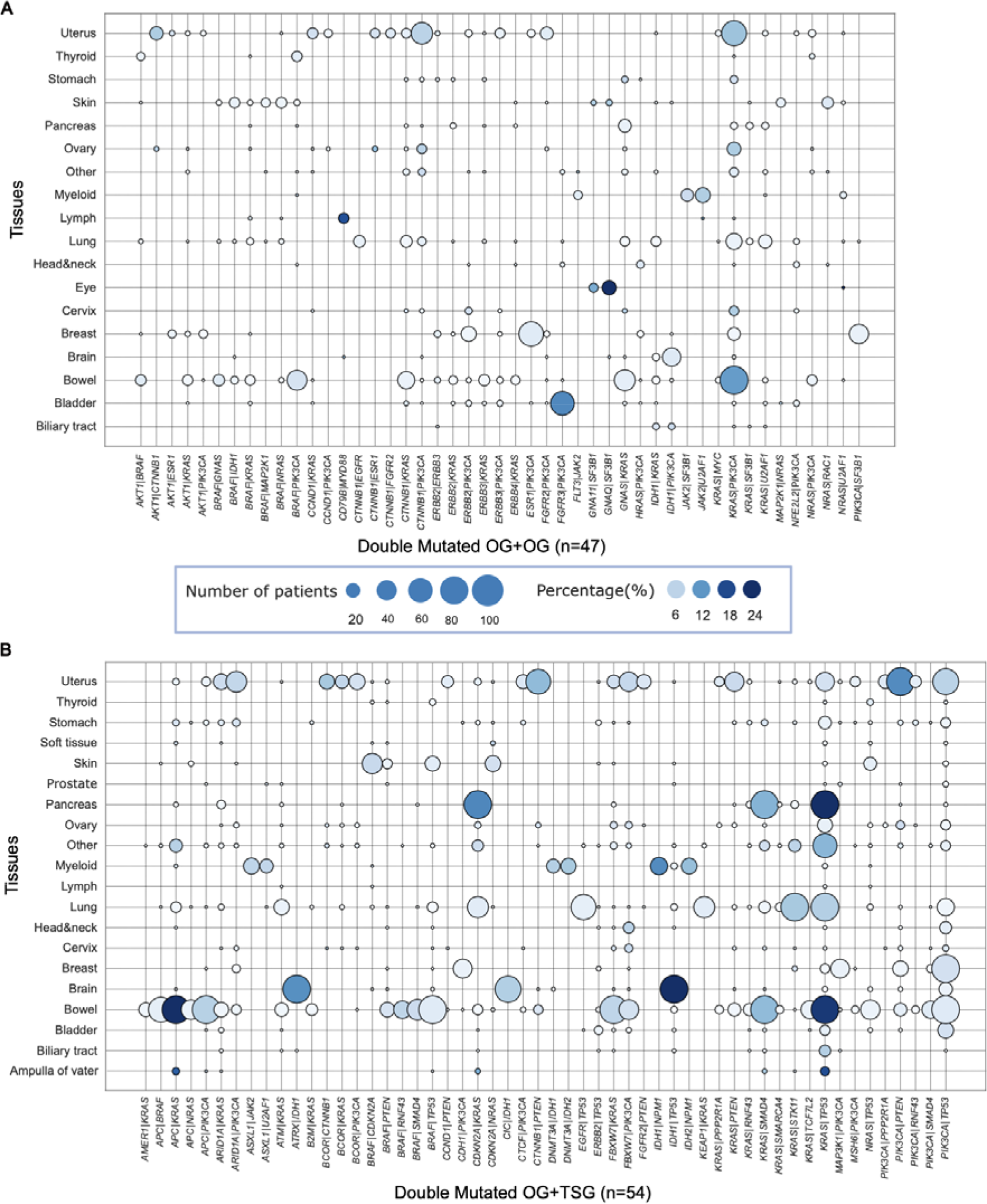
Widespread different gene double mutations in different tissues. We call a gene pair A|B if there is a double mutation where the component mutations are on the genes A and B, and the gene couple A|B is mutated on the union of tumors with double mutations on these genes. Various gene couples are prominent in different tissues. Node size is the overall number of patients with different gene double mutations in each tissue, and node color is a fraction of all double mutant tumors to all tumors in tissues. **(A)** Different oncogene pairs (mutated on at least 10 patients in a tissue) and their tissue prevalence (have mutation in at least 10 patients on an oncogene couple). *ESR1*|*PIK3CA* is significant in the breast (∼2.5%); *KRAS*|*PIK3CA* is prominent in the bowel (11%); and *CTNNB*|*PIK3CA* (∼6%) and *KRAS*|*PIK3CA* (8%) are prominent in the uterus tissues among oncogene pairs. **(B)** Different gene couples where one constituent is an oncogene and the other is a tumor suppressor gene (mutated on at least 20 patients in a tissue) and their tissue prevalence (have mutation in at least 20 patients on a gene couple). For oncogene and tumor suppressor pairs, *PIK3CA*|*TP53* (∼4%), *MAP3K1*|*PIK3CA* (1.5) and *CDH1*|*PIK3CA* (∼1.5%) are prominent in breast tissue. Similarly, the pairs *KRAS*|*SMAD4* (9%), *KRAS*|*TP53* (33%) and *CDKN2A*|*KRAS (*13%) are enriched in the pancreas tissue.

*ESR1*|*PIK3CA* oncogene pair is significant in the breast (∼2.5%); *KRAS*|*PIK3CA* is prominent in the bowel (11%); and *CTNNB*|*PIK3CA* (∼6%) and *KRAS*|*PIK3CA* (8%) are prominent in the uterus tissues. There is a wide range of oncogene and tumor suppressor pairs in bowel tissue, a well as *PIK3CA*|*TP53* (∼4%), *MAP3K1*|*PIK3CA* (1.5) and *CDH1*|*PIK3CA* (∼1.5%) in breast tissue. Along similar lines, the gene pairs *KRAS*|*SMAD4* (9%), *KRAS*|*TP53* (33%) and *CDKN2A*|*KRAS (*13%) are enriched in the pancreas tissue.

In many cases, oncogene mutations, reduce function in tumor suppressors or activating mutations in critical system components culminate in hyper-activation of the PI3Kα signaling pathway in breast cancer. Simultaneous mutations of *PIK3CA* and *MAP3K1* are found in ∼11% of *PIK3CA-* mutant tumors. Disruption of *MAP3K1* increases downstream signaling and phosphorylation of *AKT* while decreasing susceptibility to certain *AKT* and PI3Kα/δ inhibitors(73). *MAP3K1* mutational status might be a predictive biomarker for evaluating PI3K pathway inhibitor trials. The coexistence of *KRAS* and *PIK3CA* mutations in cells implies potential synergistic hyperactivation of the Ras/MAPK and PI3K/Akt oncogenic pathways. A study conducted on 655 colorectal cancer (CRC) patients indicated that co-occurring mutations in *KRAS* and *PIK3CA* were significantly associated with aggressive clinicopathological features(74). While we did not observe co-occurrence of *BRAF*/*RAS* mutations, a whole-genome study implied that *KRAS*, *NRAS*, *BRAF*, and *PIK3CA* tend to mutate concurrently, be prognostic of tumor, with possible associations of specific sex-related mutations (75). *KRAS* and *SMAD4* mutations resulted in significantly shorter relapseLfree survival in pancreatic ductal adenocarcinoma (PDAC) samples(76).

Combining the significant concomitant mutations in our dataset with results from the literature in a tissue-specific way emphasize the prognostic role of these mutations. Functional assays can be used for further evaluation of such doublets and their therapeutic roles.

### Double mutations can change transcriptional output

A driver alteration may affect the signaling pathways and the transcriptional profile of genes. In a transcriptional regulation network, these altered genes can contribute to upstream activities through feedback loops (77). We searched for transcriptional differences between tumors having double mutations in different gene pairs and their single mutation counterparts.

One of the most frequent mutations in pancreatic cancer patients is *KRAS*^G12D^. It pairs with a mutation in *TP53*, which impairs DNA binding. In the pan-cancer dataset, there are 1502 PAAD (Pancreatic adenocarcinoma) tumors, 1007 of these tumors have *KRAS*^G12D^ mutations. There are 67 significant mutation doublets composed of *KRAS*^G12^ mutations and at least one mutation in TP53 that exists in 397 patient tumors. Among them, mutations at positions 248 and 273 are directly in contact with DNA. Position 175 is far from the DNA binding region however it is in contact with a Zinc ion and a mutation at that position destabilizes p53, preventing its binding to DNA. Some tumors have significant double mutations composed of *TP53* and *KRAS*^G12D^ mutations. Because this coupling likely impacts transcriptional regulation, we compared the transcriptome profiles of Group 1 (defined as PAAD tumors having at least one significant doublet composed of *KRAS^G1^*^2D^ and *TP53* mutations) to Group 2 (PAAD tumors having mutation either in *KRAS* or in *TP53* or mutation in any of these genes that does not contribute to a significant doublet) (Figure 3A). Group 1 has 24 patient tumors. Group 2 has 71, in 35 of which TP53 is wild type in 25 samples and the rest have TP53 mutations. None contributes to co-occurring mutations. Using a conventional transcriptomic analysis, we obtained 394 differentially expressed genes between Group 1 and Group 2 in PAAD samples (Figure 3A, 137 upregulated and 257 downregulated genes where p-value < 0.01 (Mann Whitney U Test) and |log_2_FC| > 0.5). This allowed us to identify *TP53* mutations contributing to doublets in transcriptional regulation. A set of genes in immune response, positive regulation of cell proliferation and cell-cell signaling, are enriched in Group 1 compared to Group 2. We discovered significant upstream transcription factors (TFs) that regulate differentially expressed genes using the TRRUST (version 2) dataset, a manually curated database of human and mouse transcriptional regulatory networks. As a result, 55 TFs are retrieved as the main regulators including *SP1 (*encodes Sp1), *SP3* (encodes Sp3), *NFKD1*, *JUN* and *TP53* (encodes p53) (Figure 3B). We constructed the network of the TFs and KRAS with CancerGeneNet (78) and as expected, observed that the proliferation phenotype is upregulated (Figure 3C), consistent with *TP53* driver mutations significantly affecting the transcriptional output compared to the single mutant counterparts (79).

**Figure 3.**
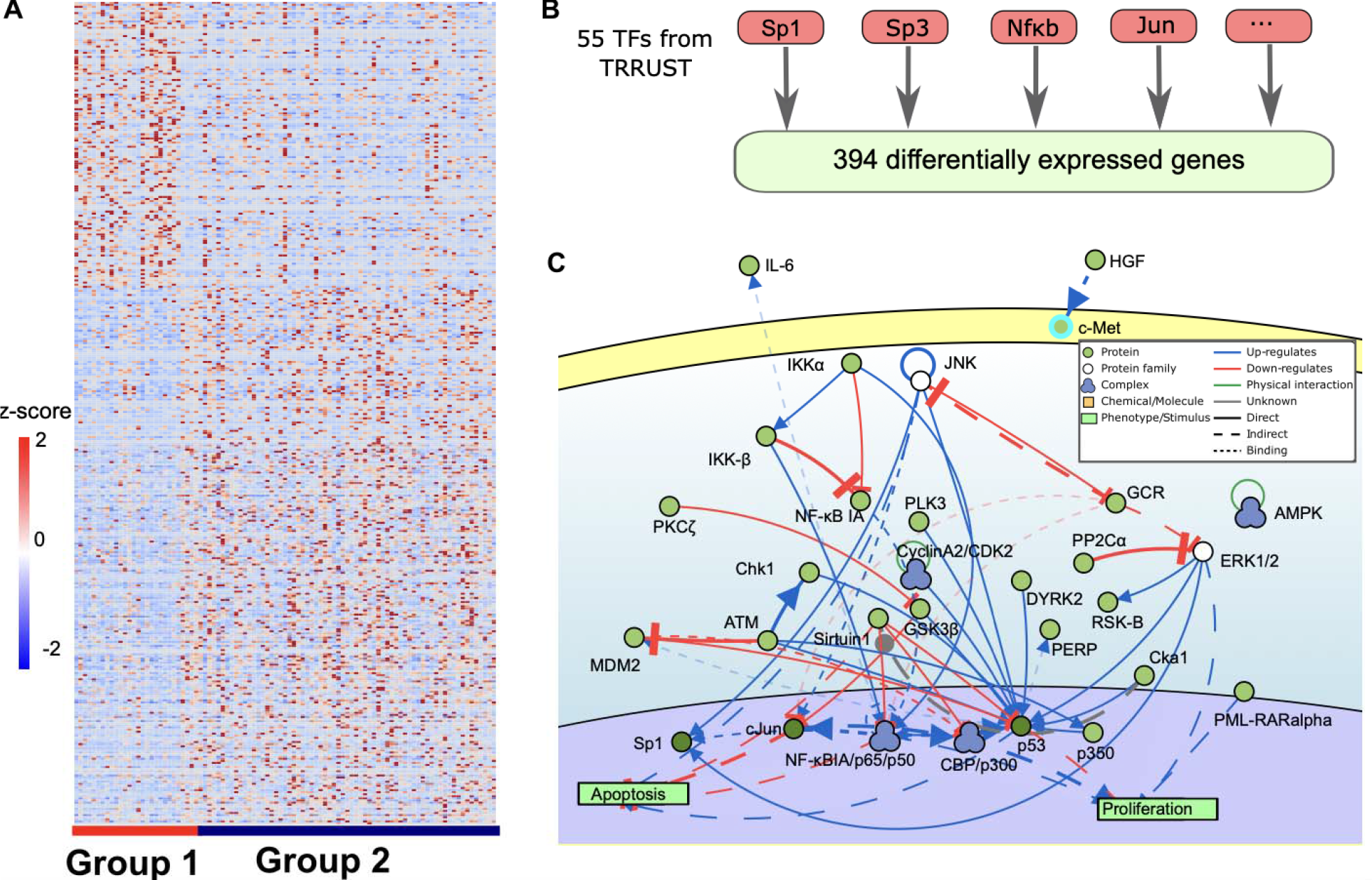
Transcriptome analysis of tumors with double mutation. **(A)** Comparison of gene expression profiles of double mutant and single mutant TP53 cases in KRAS^G12^ mutant PAAD tumors revealed 394 differentially expressed genes (151 upregulated and 243 downregulated) between Group 1 and Group 2. Group 1: PAAD tumors having at least one significant doublet composed of *KRAS^G1^*^2D^ and TP53 mutations); Group 2: PAAD tumors having mutation either in KRAS or in TP53 or mutation in any of these genes that does not contribute to a significant doublet. The heatmap shows the z-scores of differentially expressed genes (DEGs) across patient tumors. There are 137 upregulated and 257 downregulated genes. **(B)** 55 TFs are retrieved as the transcription factors of differentially expressed genes including the main regulators *SP1*, *SP3*, *NFKD1*, *JUN* and *TP53* obtained from TRRUST version 2 dataset, a manually curated database of human and mouse transcriptional regulatory networks. *SP1 (*encodes Sp1), *SP3* (encode Sp3), *NFKD1*, *JUN* and *TP53* (encodes p53). **(C)** The network of *KRAS* and important transcription factors linked to proliferation and apoptosis phenotypes was generated by CancerGeneNet^66^.

Sp1 (Specificity protein 1) is overexpressed in a variety of tumor types and is a predictor of poor patient survival. High expression levels of Sp1, Sp3, and Sp4 in cancer cells indicate that each of the three TFs contributes to the development, survival, and migration/invasion of cancer cell lines to several tissues including the pancreas (80,81). Co-occurring *TP53* and *KRAS* mutations promote tumor growth and metastasis in pancreatic ductal adenocarcinoma (PDAC) subtypes via interaction with the CREB-1 protein (cAMP responsive element binding protein 1)(82). Mutant p53 and CREB-1 upregulate *FOXA1*, a pro-metastatic pioneer TF that activates its transcriptional network while promoting WNT/β-catenin signaling, the primary driver of PDAC metastasis.

CREB-1 inhibition is a novel therapeutic strategy that aims to prevent oncogenic *KRAS* and mutant p53 from cooperating to minimize metastasis. This led to significant decrease in the expression of *FOXA1* and β-catenin, as well as reduced PDAC metastasis. In small-sized *KRAS*- transformed PDAC, *TP53* missense mutation was associated with poor tumor differentiation and revealed gain-of-function properties (83).

### Functionally equivalent alterations do not co-exist in tumors

Alterations may have similar phenotypic outcomes; we dub them ‘functionally equivalent’. According to our analysis, these alterations either rarely or never coexist in tumor tissues and appear mutually exclusive where exclusive alterations have similar signaling outputs. *KRAS*^G12D^ is coupled with alterations in a context-specific way. *KRAS*^G12D^ and the deep deletion of *CDKN2A* predominantly co-exist in pancreatic cancers. Mutually exclusive alterations may exist in different pathways but result in similar phenotypic effects. Although double mutations are extremely rare in *KRAS*, they promote a downstream alteration that leads to proliferation phenotype. Figures 4A and 4C show the presence of these mutations in *KRAS*^G12D^ mutated PAAD tumors. Among these alterations, mutations in *SMAD4*, *CDKN2A*, *U2AF1*, and *GNAS* are mutually exclusive and rarely coexist. *CDKN2A* and *SMAD4* mutations are in binding regions and rarely co-occur in tumor proteins. *CDKN2A* mutations (R58*/Q, R80*, H83N/R/Y, and D84G) are in the interface with *CDK6* (PDB: 1bi7) (Figure 4B). All, except R58*/Q, are spatially clustered. Disruption of *CDKN2A*-*CDK6* interaction inhibits their ability to interact with cyclin D and phosphorylate the retinoblastoma protein, thus down regulating proliferation. *SMAD4* mutations at D351 and R361 are in the *SMAD4*/*SMAD3* heterodimer interface(84) (PDB: 1ygs (dimer), 1u7f (trimer)) (Figure 4D). These mutations disrupt *SMAD* homo- and hetero-oligomerization. All mutually exclusive mutations shown in Figure 6 are on non-overlapping paths linked to *SMAD4* (encodes Smad4) and their functional outcome may be similar. *U2AF1* and *GNAS* (encodes Gαs)paths are linked to Ras and downstream proliferation phenotype (Figure 4E).

**Figure 4.**
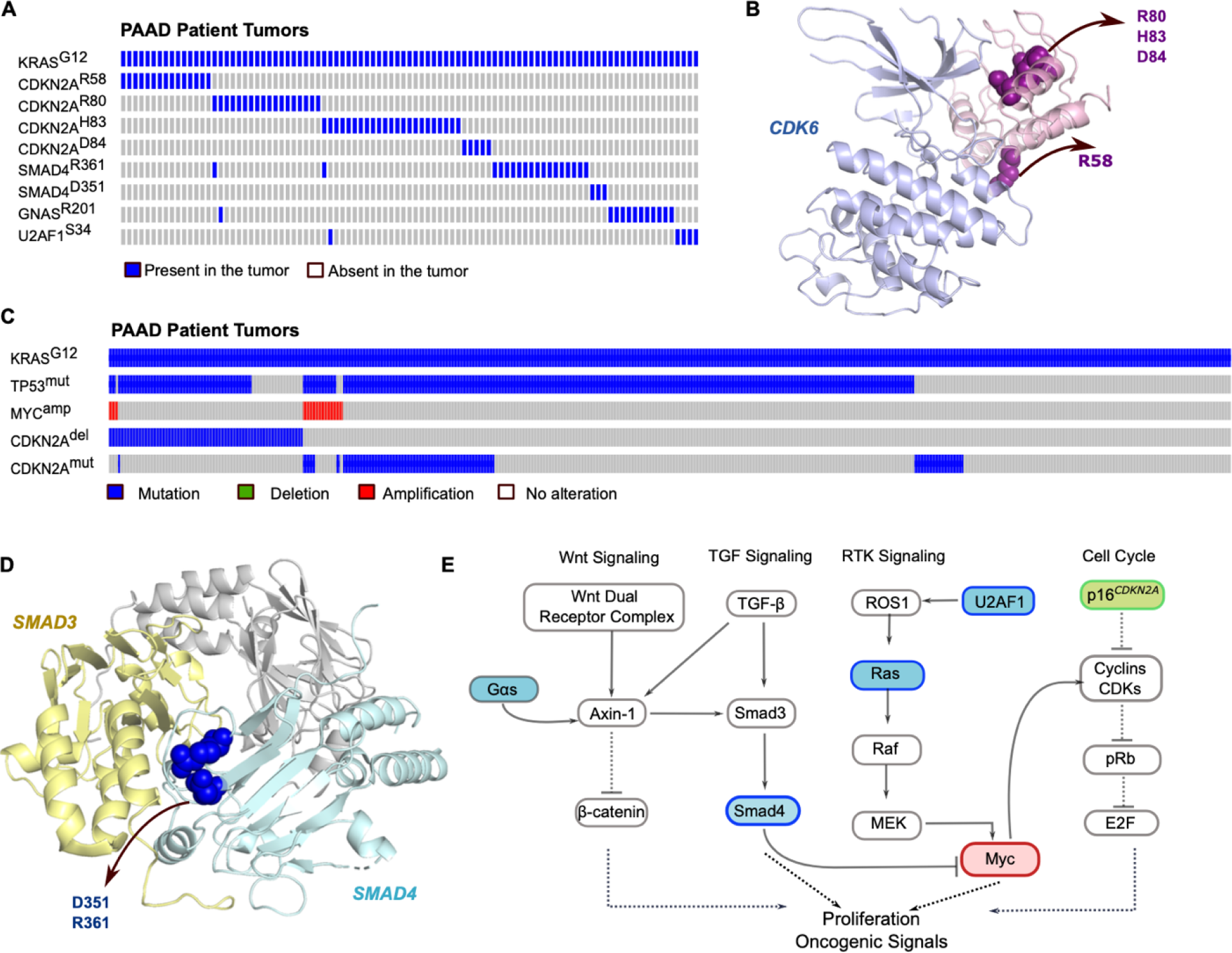
Different gene double mutations in PAAD tumors. **(A)** Oncoprint of the mutations that coexist with *KRAS*^G12^. *KRAS*^G12^ partners on *CDKN2A*, *SMAD4*, *GNAS*, and *U2AF1* shows a mutually exclusive pattern. **(B)** *CDKN2A*/*CDK6* complex. The mutations R58, R80, H83, and D84 are in the interface region between *CDKN2A* and *CDK6.* **(C)** *SMAD4* mutations D351 and R361 are in the interface between *SMAD4*/*SMAD3* complex. **(D)** Oncoprint of the most frequent alterations that coexist with *KRAS*^G12^. A *TP53* mutation on the DNA binding domain accompanies *KRAS*^G12^ and this couple is followed by either *MYC* amplification or *CDKN2A* deletion. **(E)** A subnetwork of highly frequent alterations coexisting with *KRAS*^G12^ leads to proliferation and oncogenic signals.

Co-occurrence of *MYC* amplification and *CDKN2A* deletion is extremely rare. *MYC* amplification promotes proliferation while tumor suppressor gene *CDKN2A* functions as a negative regulator of the proliferation of normal cells. They cooperate as shown by at least one *TP53* mutation in the DNA-binding domain. This is consistent with the absence of potent multiple alterations on the same, or on functionally similar pathways resulting in similar phenotype, as they can promote oncogene-induced senescence.

### Cancer type-specific subnetworks

The proportion of double mutant tumors varies across different tissues. Genes harboring double mutations are commonly in different pathways. To decrease this heterogeneity to some extent, we checked the proportion of double mutant tumors in select tissues across pathways that harbor different gene double mutation constituents, by calculating the fraction of double mutant tumors in each tissue for each of the 3424 significant different gene double mutations.

We evaluated 46 signaling pathways from KEGG. We obtained tumors with double-mutant pathways by taking the union of all double mutants when at least one of the genes belong to the corresponding pathway. We filtered the pathways that have less than 80 double mutant tumors, and tissues and pathways that have less than 10% double mutant tumor rate. We constructed a bubble plot by representing the number and fraction of pathway double-mutant tumors in each tissue as the node size and the node color, respectively (Figure 5A). On the x-axis and y-axis there are 10 tissues and 25 pathways, respectively. Bowel and uterus tissues have double mutations across all the listed pathways. On the other hand, double mutations in breast, lung and pancreas tissues accumulate in certain pathways including PI3K-Akt, MAPK, FoxO, Neurotrophin, mTOR, etc.

**Figure 5.**
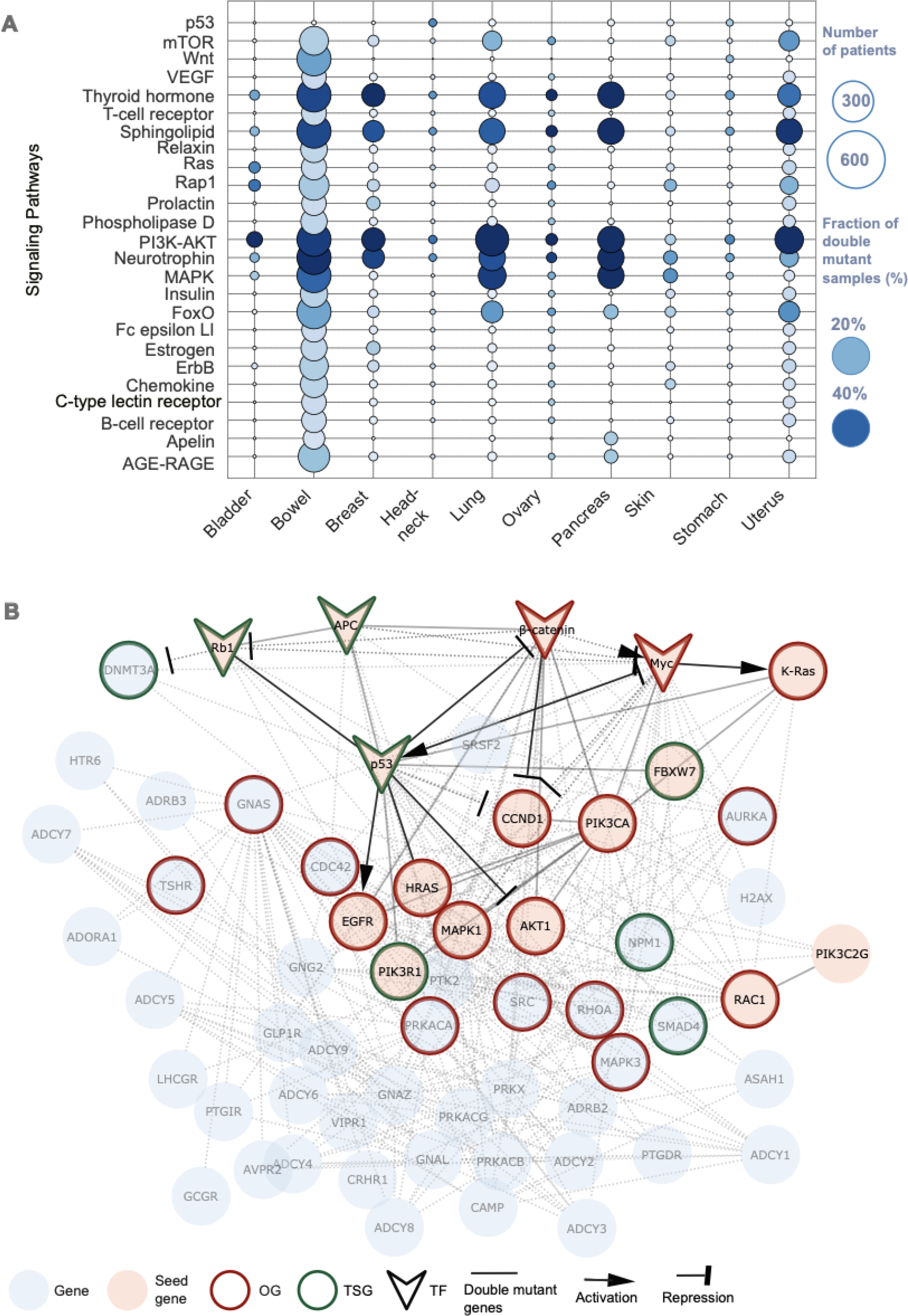
Breast cancer type specific subnetwork. (A) Bubble plot representing the number and fraction of pathway-double-mutant tumors in each tissue by the node size and the node color, respectively. On the x-axis and y-axis there are 10 tissues and 25 pathways, respectively. Bowel and uterus tissues have double mutations across all the listed pathways while double mutations in breast, lung and pancreas tissues accumulate in pathways including PI3K-AKT, MAPK, FoxO, Neurotrophin, mTOR, etc. **(B)** Breast cancer specific subnetwork obtained with the Page Rank algorithm from Omni Path with the seed genes that have double mutations and tissue specific double mutant fraction is greater than 0.5. Blue nodes are the genes coming from the PPI, pink nodes are the seed genes which are double mutation components. Transcription factors (TFs) are V-shaped nodes. Border color is green if the gene is tumor suppressor gene and red if the gene is an oncogene. The edges are solid lines if there is an edge between the nodes in the PPI and the connected nodes that contribute to a double mutation. Dashed lines depict the direct interaction in the PPI. Source and target shape are derived from TRRUST representing the activation or repression. TFs *RB1* (encodes the protein Rb1), *APC* (encodes the protein APC), *CTNNB1* (encodes Catenin Beta-1), *MYC* (encodes the protein Myc), *TP53* (encodes the protein p53).

Based on the observation that double mutations in breast, lung and pancreas tissues accumulate in specific pathways, we constructed cancer type specific subnetworks for breast and pancreatic tissues with seed genes that have double mutations, and the tissue specific double mutant fraction is greater than a certain threshold (thresholds are as follows: breast, 0.5; pancreas, 1.5).

There are 8512 tumors in breast and 1853 in pancreas tissues. We obtained a directed PPI network from OmniPath (64) with 20,029 nodes and 276,179 edges. For breast cancer, we used 15 seed genes which harbor double mutations in more than 0.5% of the breast cancer tumors. We set the initialization scores as 1 for the Page Rank algorithm with α = 0.85 and reconstructed the subnetworks for breast and pancreas cancers. There are 59 nodes and 343 edges for breast (Figure 5B) and 40 nodes and 267 edges for pancreas cancer (Figure S2). After reconstructing the cancer-type-specific subnetworks, we retrieved the list of TFs regulating the subnetwork nodes and derived the regulation information (activation or repression) for determining source and target arrow types.

For breast cancer, the resulting subnetwork includes the TFs *RB1* (encodes the protein Rb1), *APC* (encodes the protein APC), *CTNNB1* (encodes Catenin Beta-1), *MYC* (encodes the protein Myc), *TP53* (encodes the protein p53). The nodes which are oncogenes or tumor suppressor genes have red and green borders, respectively. Conducting gene set enrichment analysis in MSigDB(85,86) and Webgestalt(87) yielded that subnetwork genes enriched in the following pathways (Figure S3A-B): Signaling by GPCR (41 genes overlapped), G-alpha signaling events (28 genes), Chemokine signaling pathway (25 genes), GnRH signaling pathway (21 genes), Gap junction (20 genes), Endothelins (19 genes), LPA receptor mediated events (19 genes).

G-protein-coupled receptors (GPCRs) comprise a large family of cell-surface receptors, which have recently emerged as key players in tumorigenesis, angiogenesis and metastasis (88). Heterotrimeric G-proteins transmit mitogen-induced GPCR signals to modulate a wide range of signaling pathways. The multitude of responses elicited by GPCRs relies on the confluence of a network of transduction cascades implicated in diverse physiological processes as well as tumor development (89). GnRH (gonadotropin hormone-releasing hormone) agonists directly inhibit the growth and invasion of cancerous cells while also suppressing the production and release of ovarian steroids (90,91). Genes belonging to these pathways can be evaluated as potential drug targets for breast cancer. Through binding tumor cells chemokine receptors, chemokines produced by the tumor itself, cancer-associated fibroblasts, and infiltrating leukocytes(92,93), activate signaling pathways, such as PI3K/AKT/NF-κB and MAPK/ERK pathway, promoting cancer cell proliferation(94,95). Breast cancer is prone to chemokines and their receptors. Targeting their regulation could lead to a novel approach to breast cancer treatment (96). The TFs *APC* (encodes APC), *RB1* (encodes pRb), *DNMT3A*, *TP53* (encodes p53), *MYC*, *CTNNB1* (encodes Beta Catenin-1) and *SMAD4* (encodes Smad4) contribute to a different double mutation in the pancreas cancer subnetwork. Conducting the gene set enrichment analysis on the network nodes with Webgestalt (Figure S4A-B), we see the upregulated pathways, including PI3K-Akt, RHO GTPase signaling and their effectors and platelet activation, and aggregation.

Control of cell shape, cytokinesis, adhesion, and motility is largely dependent on the Rho family of small GTPases. Crucially, these mechanisms must be viewed within the framework of a multifaceted three-dimensional (3D) environment, wherein the tumor cells and the host environment engage in mutual feedback and cross-talk (97). Rho GTPases are essential for the development of dynamic actin-rich membrane protrusions as well as the turnover of cell-cell and cell-extracellular matrix adhesions (97). They also facilitate pancreatic ductal cell motility and invasion. Together, our functional and genomic profiles demonstrate that the GTP-binding Rho family proteins are essential to the fatal disease’s invasive characteristic(98,99). Platelets reportedly contribute to the growth, invasion, and metastasis of various tumors, including pancreatic cancer (100,101).

Given the results of pathway enrichment analysis along with the available literature indicates that a thorough examination of the reconstructed breast cancer and pancreas cancer subnetworks from the genes carrying double mutations can offer valuable insights into the development of certain cancer types as well as targets for therapeutic intervention.

### Double mutations are mostly present in primary tumors and some rare doubles are signature of metastatic tumors

The early origins of metastatic lineages, role of driver mutations, and the nonlinear patterns of tumor progression are key long-standing questions(102). Primary and metastatic tumor tissues of certain cancer types were compared to obtain their genomic heterogeneity and homogeneity(103). Compared with primary cancers, metastatic cancers only have a moderate increase in tumor mutation burden (TMB) (55). How then the metastatic phenotypes arise at the genomic level, what are the molecular mechanisms that determine how specific oncogenic drivers interact with the related physiological programs, and what triggers their activation in support of metastasis are challenging questions (104,105). A study of 40,979 primary and metastatic tumors across 25 distinct cancer types in the AACR GENIE dataset revealed that the most commonly mutated genes among metastatic tumors were *TP53*, *MYC*, and *CDKN2A*. No discernible pattern linked to metastatic disease was identified, and of note, the driver mutation profiles were also very similar (106).

The AACR GENIE cohort has samples from primary and metastatic tumors. Having metastatic samples enabled us to investigate co-occurrence patterns of mutations enriched in metastatic and primary breast cancer. We observed that more than 70% of all double mutations exist in primary tumors. However, there is a small set of double mutations that are specific to metastatic samples. We applied the frequency pattern growth tree approach (see Methods) to obtain the association rules of molecular alterations in metastatic tumors among (Figure 6). The Metastatic Breast Cohort consists of 2385 samples, and the subtype distribution can simply be classified as 52% IDC (invasive ductal carcinoma), 25% BRCA (breast cancer), 9% ILC (invasive lobular carcinoma), 4.5% BRCNOS (breast invasive carcinoma), and the rest are other subtypes. The input matrix of the FP (Frequency Pattern) growth tree is composed of 456 mutations and 2385 metastatic breast tumors. There are 31 nodes and 51 edges in the FP growth tree of breast cancer metastatic tumors. Out of the 51 pairs in the FP growth tree, 46 (∼90%) are among the different gene double mutations which we evaluate as metastatic markers (Supplementary Table 2). All the pairs with q<0.3 with at least three double mutant tumors are co-occurring. The frequency patterns identify a strong association between mutations in *ESR1*, *GATA3*, and *PIK3CA* genes in metastatic tumors. *ESR1* mutations at positions 380, 536, 537 and 538, which are sequence neighbors, are exclusively paired with the major drivers of *PIK3CA* at positions 542, 545, and 1047 (Figure 6). In Figure 7A, the volcano plot shows the co-occurrence and mutual exclusivity tendencies of the metastatic markers in breast cancer. On the x-axis there is log10(Odds Ratio) against -log10 Fisher’s p-value on the y-axis where non-significant pairs (σ > 0.05, |Odds Ratio| > 0.75) are in gray. We also calculated the propensity of double mutations in *ESR1* and *PIK3CA* to be in metastatic tumors and doublets of each mutation with mutations in other genes. The result strongly supports mutation pairs, one from *ESR1* and one from *PIK3CA* as markers of metastatic tumors (Figure 7B, p-value < 10^-13^). *GATA3* mutations at positions 293, 335, 408 are coupled with *ESR1* or *PIK3CA* mutations. *GATA3* is also functionally related and cooperates with *ESR1* in transcriptional regulation (107). Figure 7C, evaluates all possible relationships of the mutations at the FP growth tree in the bubble plot where larger nodes indicate higher co-mutation tendency. We filtered the 330 pairs with less than three double mutant tumors.

**Figure 6.**
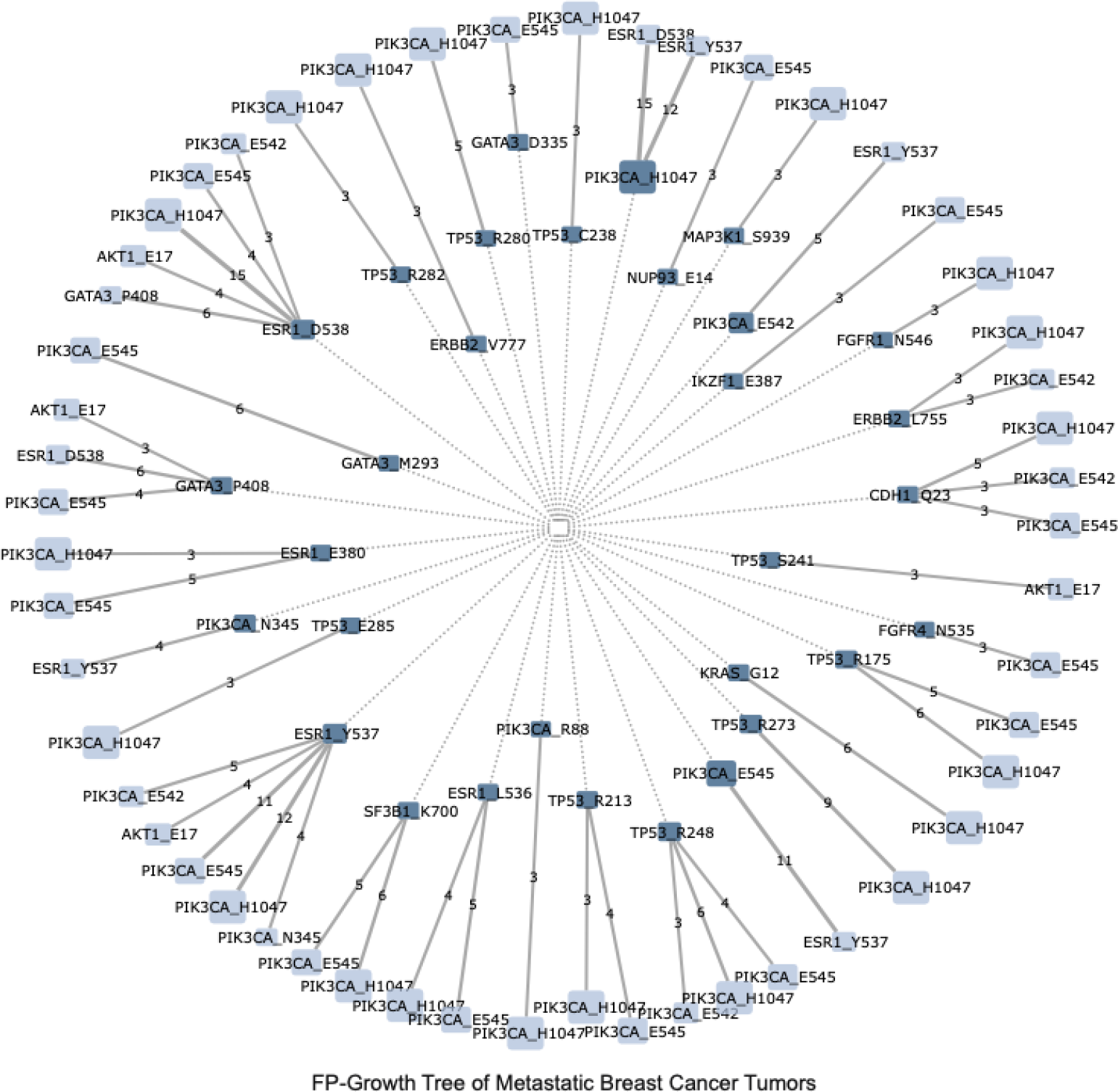
Some rare double mutations are specific to metastatic tumors. Frequency pattern growth tree of missense mutations in breast cancer metastatic tumors where links represent association between mutations. Edges represent correlation strength. Ancestral mutations are shown as dark lilac nodes, leading to consequential light lilac mutations. Mutation frequencies are represented by node sizes. Different gene double mutations in *ESR1*, *GATA3*, and *PIK3CA* genes are enriched in metastatic tumors. There are 31 nodes and 51 edges in the FP growth tree out of which 46 pairs are among the different gene double mutations which are evaluated a metastatic markers.

**Figure 7.**
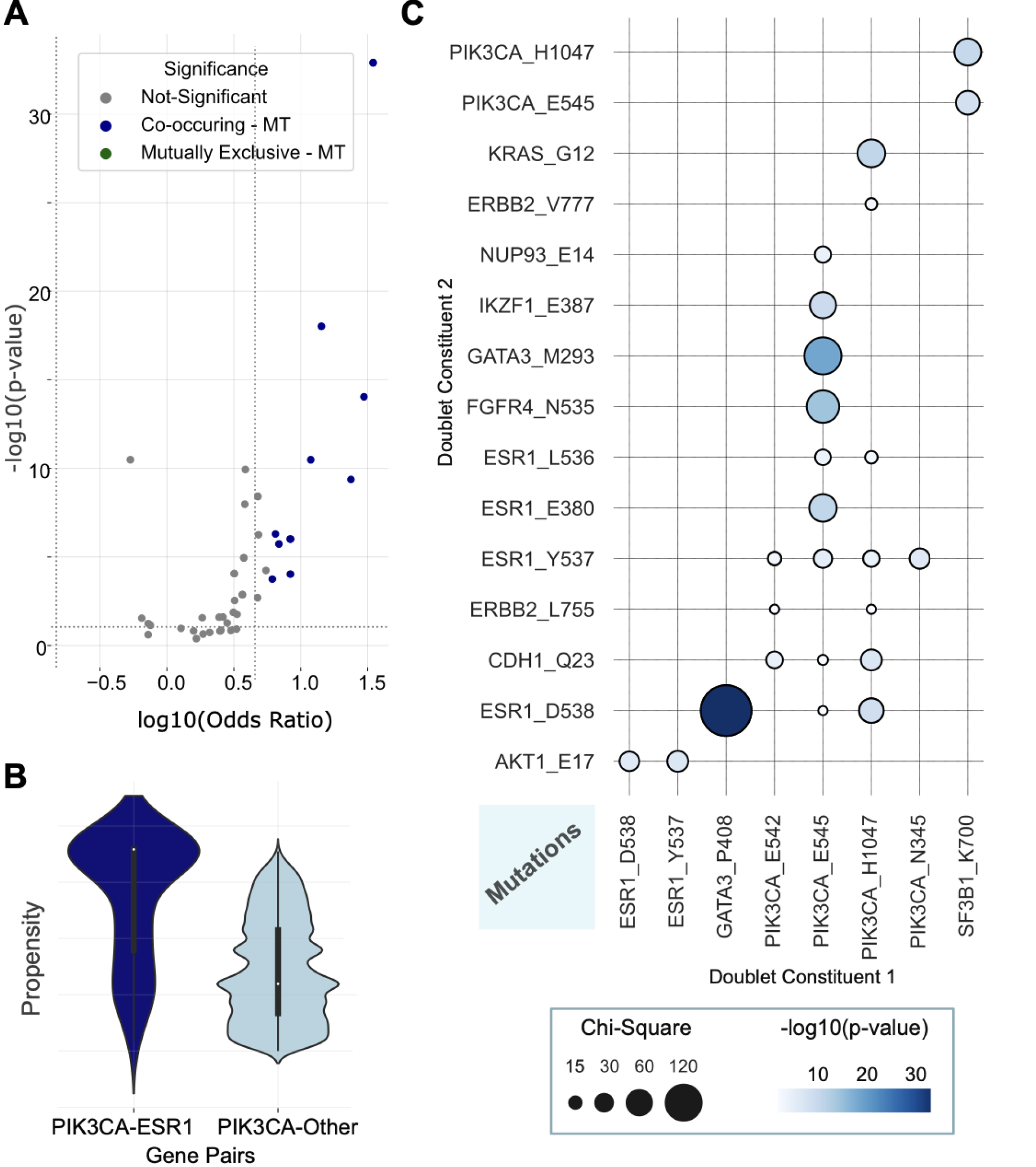
Metastatic marker statistics. (A) The Volcano plot of pairs of same cohort, log10 Odds Ratio against -log10 Fisher’s p-value, with non-significant pairs (σ > 0.05, |Odds Ratio| > 0.75) in gray. **(B)** Propensity of double mutations from *ESR1*-*PIK3CA* pair and from other genes where one side is from *ESR1* or *PIK3CA* in metastatic tumors. **(C)** The bubble plot evaluating all possible relationships of the mutations at the FP growth tree, where node color depicts coexistence characteristics, and node radius is proportional to the Chi-Square value of the same cohort.

In the supplementary data of the study(108), there is a list of actionable mutations for 1106 metastatic tumors from the Hartwig Medical Foundation (HMF) data set. Among 393 metastatic breast tumors, 22 harbor one *PIK3CA* and one *ESR1* mutation. The mutations at positions 542, 545, H1047 of *PIK3CA* confirm occurrence of a double mutation with one of the mutations at positions 380, 536, 537 and 538 on ESR1, and determined as metastatic markers by the FP growth tree algorithm.

In addition to these, we provide a tree of double, triple, and quadruple co-occurrence patterns across the pan-cancer metastatic tumors (Figure S5A-B). In the tree, the node *TP53*^R273^ is followed by either one of the mutations on ATRX at the positions 1045, 1049, and 1426 where the third node of the triplet is *IDH1*^R132^. *IDH1*, *TP53*, and *ATRX* mutations associated with low-grade astrocytomas collaborate in order to inhibit *SOX2* (a nucleosome-binding pioneer TF acting to make genes accessible to the transcription machinery) and prevent the development of human neural stem cells (109,110). Quadruples include mutations from *BRAF, MSH3, BLM, RNF43, PLK2, NBN*.

The genomic differences/similarities between primary and metastatic tumors largely remain uncharacterized and the most altered genes and in most cases the driver mutations are similar(106). Finding double or higher order interactions between the mutations in primary and metastatic cancer may offer a more comprehensive comparison between these two groups and aid in the identification of the molecular signatures that facilitate the development of multiple drug combinations targeting metastatic cancers.

## Discussion

Recent studies undertook exploration of the role of double/multiple mutations in *cis*, i.e. in the same gene (72,79,111–113). They observed that *cis* mutations increase oncogenic activity, thus cell lines and xenografts that harbor such mutations become more sensitive to certain drugs. Double mutations promote tumor growth and hyperactivate PI3K signaling in model systems. Preliminary analysis of clinical trial data indicates that double-mutant breast cancers respond better to PI3K inhibitors than single-mutant cancers (113). Inverse effects were also observed as in the case for EGFR^L858/T790^ cell line which becomes resistant to the inhibitors compared to the EGFR^L858^.

Here, we carry out analysis of co-occurring mutations, *in trans,* to identify the mutations that can cooperate to enhance the oncogenic signal. In some cases, pairs are mutually exclusive. Presumably, sporadic emergence of these pairs led to unsustainable hyperactivity of the cancer cell through signaling overdrive, either via e.g., (1) same- or redundant-pathway hotspot mutations, or (2) through a combination of a pathway mutation and mutation in a pioneer TF, such as FOXA1, which displaces linker histone H1, thereby keeping enhancer nucleosomes accessible leading to overexpression (114), and OIS, thus absent in our dataset. We identified 3424 significant gene double mutations, composed of mostly one rare and one frequent mutation. 3307 doublets are co-occurring while the rest are mutually exclusive. Frequent mutations have been cataloged as driver mutations. Even though some driver mutations are cancer-wide, coupling with another (driver or passenger) mutation increases their cancer-type specificity. We further adopted the FP-growth tree method (61), which has been developed for database systems, to find the association rules between single point mutations. Our analysis revealed association across metastatic tumors and discovered that *PIK3CA* and *ESR1* mutations co-occur in metastatic tumors and the doublets in these genes are more frequent in metastatic tumors compared to primary tumors.

Considering the additivity of the mutational impact, mutations affecting the same pathway might be mutually exclusive in a tumor to prevent functional redundancy, and especially, if strong, synthetic lethality (6,36,115), or OIS (7,21), which clarifies its molecular basis. We also observe some cases in protein interfaces, where double hotspot mutations in the same interface are exclusive, e.g., *CDKN2A* R58, R80, H83 and R84. Driver mutations co-occurrence (DCO) in cell-type specific networks can be helpful in identifying synergy and drug response (116) and identifying druggable mutations across cancer types can enhance targeted therapies (117). Some of the *in trans* double mutations that are prominent in specific tissues or cancer subtypes are associated with poor overall survival. For example, co-occurring mutations in *IDH2*/*NPM1*, *IDH1*/*NPM1*, *DNMT3A/NPM1* pairs are significantly prevalent in AML tumors which have been recently associated with poorer overall survival (118). Another such combination is mutated *RNF43* together with *BRAF*^V600E^ which significantly co-occur in our dataset. This cooperativity was defined as a marker of aggressive right-sided colorectal cancer (RCRC) subtype (119). Co-occurring mutations in *PIK3CA* and *ARID1A* in uterine endometrioid carcinoma (UEC) subtypes raise in sensitivity to PI3K inhibitors (120).

Over 70% of all double mutations exist in primary tumors; however, there is a small set of double mutations that are specific to metastatic samples. Estrogen Receptor 1 (*ESR1*) and *PIK3CA* top the list. They are components of the well-established insulin– phosphoinositide 3- kinase (PI3K) signaling cascade, known to play key roles in cancer cells(121). *AKT* is in the PI3K/mTOR pathway. *ESR1* and PI3Kα lipid kinase coordinates the glucose metabolism upstream, and *AKT* and mTOR, are protein kinases downstream of PI3K. The pathway feeds into the cell cycle, acting dominantly in protein synthesis and is a driver of malignant transformation. Also ranking high is GATA3, a pioneer transcription factor (TF)(122) controlling multiple genes in embryonic development known to be a marker for some cancers, particularly those related with the breast. *GATA3* acts upstream of FOXA1, also a pioneer TF, in mediating *ESR1*(107). It is also associated with PI3K/mTOR. Consistently, we observe *GATA3*/*ESR1* and *GATA3*/*PIK3CA* signatures. Additional candidates that we observe are *TP53*, *ERBB2* (a receptor tyrosine kinase with intrinsic tyrosine kinase activity, established to mediate breast cancer) and CDH1 (Cadherin-1 or Epithelial cadherin). These confirm and link to established major mechanisms in proliferative cancers.

Collectively, here we took up two fundamental aims in cancer research, which despite their significance to date have not been addressed: (i) discovering metastatic markers; (ii) mutations whose combinations can encode oncogene-induced senescence, thus are rare, or missing in genomic cancer databases. As to OIS (ii), we conjectured that the reason that they are non-existent is that the signal that they produce is too strong for the cell to bear. Mutations occur sporadically during the cancer evolution. Cells do not have a mechanism for an *a priori* negative selection. Rather, negative selection, or exclusion, is the outcome of cell death. Here we provide a list, a rationale, and suggest that OIS can provide the molecular basis to synthetic lethality, which has been investigated on the gene level. As to (i), to our knowledge, we provide a first candidate list of metastatic markers specific to metastatic samples, including actionable mutations for metastatic tumors, as well as a tree of double, triple, and quadruple co-occurrence patterns across the pan-cancer metastatic. This list (Supplementary Table 2) may help in early identification. Advanced computational approaches allow detection of patterns. We believe that the key points are the questions that are asked, and the interpretation. Co-occurring mutational signatures for metastatic tumors, coupled with identification of the respective proteins and pathways, can create interactive maps. Linking them with drugs can create immensely useful tools for the attending physicians (123,124).

Finally, while the usefulness of metastatic markers to drug discovery is self-evident, exactly how identification of OIS mutations in protein pairs is less so, as indeed is also the case in synthetic lethality (125). OIS restriction is observed in same- and redundant-pathways. Classically, drugs aim to block protein activity and downstream signaling. Introducing drugs that recreate their action is expected to lead to cell death. At the same time, controlling the signal is challenging, as strong signals can be associated with cell proliferation, but too strong a signal may result in oncogene-induced senescence.

## Data and code availability

The results shown here are in whole or part based upon data generated by the TCGA Research Network: https://www.cancer.gov/tcga. All the data and codes are available at https://github.com/bengiruken/DifferentGeneDoubleMutations. The codes and data for obtaining metastatic markers with the FP growth tree is available at https://github.com/ugur0sahin/FMPSeeker.

## List of abbreviations

AACR: American Association for Cancer Research
TCGA: The Cancer Genome Atlas
GENIE: Genomics Evidence Neoplasia Information Exchange
PDB: Protein Data Bank

## References

1. Boca, S.M., Kinzler, K.W., Velculescu, V.E., Vogelstein, B. and Parmigiani, G. (2010) Patient-oriented gene set analysis for cancer mutation data. Genome Biol., 11, R112.

2. Reiter, J.G., Baretti, M., Gerold, J.M., Makohon-Moore, A.P., Daud, A., Iacobuzio-Donahue, C.A., Azad, N.S., Kinzler, K.W., Nowak, M.A. and Vogelstein, B. (2019) An analysis of genetic heterogeneity in untreated cancers. Nat Rev Cancer, 19, 639–650.

3. Campbell, P.J. (2017) Cliques and Schisms of Cancer Genes. Cancer Cell, 32, 129–130.

4. Nussinov, R., Tsai, C.-J. and Jang, H. (2020) Are Parallel Proliferation Pathways Redundant? Trends Biochem. Sci., 45, 554–563.

5. Nussinov, R., Zhang, M., Maloney, R. and Jang, H. (2021) Drugging multiple same-allele driver mutations in cancer. Expert Opin. Drug Discov., 16, 823–828.

6. 6. van de Haar, J., Canisius, S., Yu, M.K., Voest, E.E., Wessels, L.F.A. and Ideker, T. (2019) Identifying Epistasis in Cancer Genomes: A Delicate Affair. Cell, 177, 1375–1383.

7. Nussinov, R., Tsai, C.-J. and Jang, H. (2022) A New View of Activating Mutations in Cancer. Cancer Res., 82, 4114–4123.

8. Nussinov, R., Tsai, C.-J. and Jang, H. (2022) Allostery, and how to define and measure signal transduction. Biophys. Chem., 283, 106766.

9. Cisowski, J., Sayin, V.I., Liu, M., Karlsson, C. and Bergo, M.O. (2016) Oncogene-induced senescence underlies the mutual exclusive nature of oncogenic KRAS and BRAF. Oncogene, 35, 1328–1333.

10. Cavalli, G., Biavasco, R., Borgiani, B. and Dagna, L. (2014) Oncogene-induced senescence as a new mechanism of disease: the paradigm of erdheim-chester disease. Front. Immunol., 5, 281.

11. Zhang, M., Jang, H. and Nussinov, R. (2021) PI3K Driver Mutations: A Biophysical Membrane-Centric Perspective. Cancer Res., 81, 237–247.

12. Nordling, C.O. (1953) A new theory on cancer-inducing mechanism. Br. J. Cancer, 7, 68–72.

13. Ashley, D.J. (1969) The two “hit” and multiple “hit” theories of carcinogenesis. Br. J. Cancer, 23, 313–328.

14. Knudson, A.G., Jr. (1971) Mutation and cancer: statistical study of retinoblastoma. Proc. Natl. Acad. Sci. U. S. A., 68, 820–823.

15. Boland, C.R. and Ricciardiello, L. (1999) How many mutations does it take to make a tumor? Proc. Natl. Acad. Sci. U. S. A., 96, 14675–14677.

16. Tomlinson, I., Sasieni, P. and Bodmer, W. (2002) How many mutations in a cancer? Am. J. Pathol., 160, 755–758.

17. Tomasetti, C., Marchionni, L., Nowak, M.A., Parmigiani, G. and Vogelstein, B. (2015) Only three driver gene mutations are required for the development of lung and colorectal cancers. Proc. Natl. Acad. Sci. U. S. A., 112, 118–123.

18. Nussinov, R., Jang, H., Tsai, C.-J. and Cheng, F. (2019) Precision medicine review: rare driver mutations and their biophysical classification. Biophys. Rev., 11, 5–19.

19. 19. Ahronian, L.G., Sennott, E.M., Van Allen, E.M., Wagle, N., Kwak, E.L., Faris, J.E., Godfrey, J.T., Nishimura, K., Lynch, K.D., Mermel, C.H., et al. (2015) Clinical Acquired Resistance to RAF Inhibitor Combinations in BRAF-Mutant Colorectal Cancer through MAPK Pathway Alterations. Cancer Discov., 5, 358–367.

20. Rajagopalan, H., Bardelli, A., Lengauer, C., Kinzler, K.W., Vogelstein, B. and Velculescu, V.E. (2002) Tumorigenesis: RAF/RAS oncogenes and mismatch-repair status. Nature, 418, 934.

21. Cisowski, J. and Bergo, M.O. (2017) What makes oncogenes mutually exclusive? Small GTPases, 8, 187–192.

22. Schmitt, C.A., Wang, B. and Demaria, M. (2022) Senescence and cancer — role and therapeutic opportunities. Nat. Rev. Clin. Oncol., 19, 619–636.

23. Wang, B., Kohli, J. and Demaria, M. (2020) Senescent Cells in Cancer Therapy: Friends or Foes? Trends Cancer Res., 6, 838–857.

24. Liu, X.-L., Ding, J. and Meng, L.-H. (2018) Oncogene-induced senescence: a double edged sword in cancer. Acta Pharmacol. Sin., 39, 1553–1558.

25. Zhu, H., Blake, S., Kusuma, F.K., Pearson, R.B., Kang, J. and Chan, K.T. (2020) Oncogene-induced senescence: From biology to therapy. Mech. Ageing Dev., 187, 111229.

26. Jung, S.H., Hwang, H.J., Kang, D., Park, H.A., Lee, H.C., Jeong, D., Lee, K., Park, H.J., Ko, Y.-G. and Lee, J.-S. (2019) mTOR kinase leads to PTEN-loss-induced cellular senescence by phosphorylating p53. Oncogene, 38, 1639–1650.

27. Chan, K.T., Blake, S., Zhu, H., Kang, J., Trigos, A.S., Madhamshettiwar, P.B., Diesch, J., Paavolainen, L., Horvath, P., Hannan, R.D. et al. (2020) A functional genetic screen defines the AKT-induced senescence signaling network. Cell Death Differ., 27, 725–741.

28. Tu, Z., Aird, K.M. and Zhang, R. (2012) RAS, cellular senescence and transformation: the BRCA1 DNA repair pathway at the crossroads. Small GTPases, 3, 163–167.

29. Deng, Y., Luo, S., Deng, C., Luo, T., Yin, W., Zhang, H., Zhang, Y., Zhang, X., Lan, Y., Ping, Y. et al. (2019) Identifying mutual exclusivity across cancer genomes: computational approaches to discover genetic interaction and reveal tumor vulnerability. Brief. Bioinform., 20, 254–266.

30. Blair, L.M., Juan, J.M., Sebastian, L., Tran, V.B., Nie, W., Wall, G.D., Gerceker, M., Lai, I.K., Apilado, E.A., Grenot, G. et al. (2023) Oncogenic context shapes the fitness landscape of tumor suppression. Nat. Commun., 14, 6422.

31. Park, S., Supek, F. and Lehner, B. (2021) Higher order genetic interactions switch cancer genes from two-hit to one-hit drivers. Nat. Commun., 12, 1–10.

32. Sinkala, M. (2023) Mutational landscape of cancer-driver genes across human cancers. Sci Rep, 13, 12742.

33. Tokheim, C. and Karchin, R. (2019) CHASMplus Reveals the Scope of Somatic Missense Mutations Driving Human Cancers. Cell Syst, 9, 9–23 e28.

34. Rogers, Z.N., McFarland, C.D., Winters, I.P., Naranjo, S., Chuang, C.-H., Petrov, D. and Winslow, M.M. (2017) A quantitative and multiplexed approach to uncover the fitness landscape of tumor suppression in vivo. Nat. Methods, 14, 737–742.

35. Harris, T.J.R. and McCormick, F. (2010) The molecular pathology of cancer. Nat. Rev. Clin. Oncol., 7, 251–265.

36. Sanchez-Vega, F., Mina, M., Armenia, J., Chatila, W.K., Luna, A., La, K.C., Dimitriadoy, S., Liu, D.L., Kantheti, H.S., Saghafinia, S. et al. (2018) Oncogenic Signaling Pathways in The Cancer Genome Atlas. Cell, 173, 321–337.e310.

37. Liu, S., Liu, J., Xie, Y., Zhai, T., Hinderer, E.W., Stromberg, A.J., Vanderford, N.L., Kolesar, J.M., Moseley, H.N.B., Chen, L. et al. (2021) MEScan: a powerful statistical framework for genome-scale mutual exclusivity analysis of cancer mutations. Bioinformatics, 37, 1189–1197.

38. Hua, X., Hyland, P.L., Huang, J., Song, L., Zhu, B., Caporaso, N.E., Landi, M.T., Chatterjee, N. and Shi, J. (2016) MEGSA: A Powerful and Flexible Framework for Analyzing Mutual Exclusivity of Tumor Mutations. Am. J. Hum. Genet., 98, 442–455.

39. Leiserson, M.D.M., Wu, H.-T., Vandin, F. and Raphael, B.J. (2015) CoMEt: a statistical approach to identify combinations of mutually exclusive alterations in cancer. Genome Biol., 16, 160.

40. Ciriello, G., Cerami, E., Aksoy, B.A., Sander, C. and Schultz, N. (2013) Using MEMo to discover mutual exclusivity modules in cancer. *Curr. Protoc. Bioinformatics*, **Chapter** 8, 8.17.11-18.17.12.

41. Zhang, J., Wu, L.-Y., Zhang, X.-S. and Zhang, S. (2014) Discovery of co-occurring driver pathways in cancer. BMC Bioinformatics, 15, 271.

42. Mina, M., Raynaud, F., Tavernari, D., Battistello, E., Sungalee, S., Saghafinia, S., Laessle, T., Sanchez-Vega, F., Schultz, N., Oricchio, E. and Ciriello, G. (2017) Conditional Selection of Genomic Alterations Dictates Cancer Evolution and Oncogenic Dependencies. Cancer Cell, 32, 155–168.e156.

43. Jubb, H.C., Pandurangan, A.P., Turner, M.A., Ochoa-Montaño, B., Blundell, T.L. and Ascher, D.B. (2017) Mutations at protein-protein interfaces: Small changes over big surfaces have large impacts on human health. Prog. Biophys. Mol. Biol., 128, 3–13.

44. Cheng, F., Zhao, J., Wang, Y., Lu, W., Liu, Z., Zhou, Y., Martin, W.R., Wang, R., Huang, J., Hao, T. et al. (2021) Comprehensive characterization of protein-protein interactions perturbed by disease mutations. Nat. Genet., 53, 342–353.

45. Cheng, F., Liang, H., Butte, A.J., Eng, C. and Nussinov, R. (2019) Personal Mutanomes Meet Modern Oncology Drug Discovery and Precision Health. Pharmacol. Rev., 71, 1–19.

46. Nussinov, R., Jang, H., Tsai, C.-J. and Cheng, F. (2019) Review: Precision medicine and driver mutations: Computational methods, functional assays and conformational principles for interpreting cancer drivers. PLoS Comput. Biol., 15, e1006658.

47. Ivanov, A.A. (2020) Explore Protein-Protein Interactions for Cancer Target Discovery Using the OncoPPi Portal. Methods Mol. Biol., 2074, 145–164.

48. Ivanov, A.A., Revennaugh, B., Rusnak, L., Gonzalez-Pecchi, V., Mo, X., Johns, M.A., Du, Y., Cooper, L.A.D., Moreno, C.S., Khuri, F.R. and Fu, H. (2018) The OncoPPi Portal: an integrative resource to explore and prioritize protein-protein interactions for cancer target discovery. Bioinformatics, 34, 1183–1191.

49. Nussinov, R., Tsai, C.-J. and Jang, H. (2021) Anticancer drug resistance: An update and perspective. Drug Resist. Updat., 59, 100796.

50. Chen, C., Shi, C., Huang, X., Zheng, J., Zhu, Z., Li, Q., Qiu, S., Huang, Z., Zhuang, Z., Wu, R. et al. (2019) Molecular Profiles and Metastasis Markers in Chinese Patients with Gastric Carcinoma. Sci. Rep., 9, 13995.

51. Zhang, Y., Chen, Y., Yang, C., Seger, N., Hesla, A.C., Tsagkozis, P., Larsson, O., Lin, Y. and Haglund, F. (2021) TERT promoter mutation is an objective clinical marker for disease progression in chondrosarcoma. Mod. Pathol., 34, 2020–2027.

52. Liu, R., Rizzo, S., Waliany, S., Garmhausen, M.R., Pal, N., Huang, Z., Chaudhary, N., Wang, L., Harbron, C., Neal, J. et al. (2022) Systematic pan-cancer analysis of mutation- treatment interactions using large real-world clinicogenomics data. Nat. Med., 28, 1656–1661.

53. Smith, M.R., Wang, Y., D’Agostino, R., Jr., Liu, Y., Ruiz, J., Lycan, T., Oliver, G., Miller, L.D., Topaloglu, U., Pinkney, J. et al. (2023) Prognostic Mutational Signatures of NSCLC Patients treated with chemotherapy, immunotherapy and chemoimmunotherapy. NPJ Precis Oncol, 7, 34.

54. Zhang, F., Wang, J., Xu, Y., Cai, S., Li, T., Wang, G., Li, C., Zhao, L. and Hu, Y. (2022) Co-occurring genomic alterations and immunotherapy efficacy in NSCLC. NPJ Precis Oncol, 6, 4.

55. 55. Martínez-Jiménez, F., Movasati, A., Brunner, S.R., Nguyen, L., Priestley, P., Cuppen, E. and Van Hoeck, A. (2023) Pan-cancer whole-genome comparison of primary and metastatic solid tumours. Nature, 618, 333–341.

56. Cerami, E., Gao, J., Dogrusoz, U., Gross, B.E., Sumer, S.O., Aksoy, B.A., Jacobsen, A., Byrne, C.J., Heuer, M.L., Larsson, E. et al. (2012) The cBio cancer genomics portal: an open platform for exploring multidimensional cancer genomics data. Cancer Discov., 2, 401–404.

57. Consortium, A.P.G. (2017) AACR Project GENIE: Powering Precision Medicine through an International Consortium. Cancer Discov., 7, 818–831.

58. Tamborero, D., Rubio-Perez, C., Deu-Pons, J., Schroeder, M.P., Vivancos, A., Rovira, A., Tusquets, I., Albanell, J., Rodon, J., Tabernero, J. et al. (2018) Cancer Genome Interpreter annotates the biological and clinical relevance of tumor alterations. Genome Med., 10, 25.

59. Kanehisa, M. and Goto, S. (2000) KEGG: kyoto encyclopedia of genes and genomes. Nucleic Acids Res, 28, 27–30.

60. Gu, Z. and Hubschmann, D. (2022) Make Interactive Complex Heatmaps in R. Bioinformatics, 38, 1460–1462.

61. 61. Jiawei Han School of Computing Science, S.F.U., Jian Pei School of Computing Science, S.F.U. and Yiwen Yin School of Computing Science, S.F.U. (2000) Mining frequent patterns without candidate generation. *ACM SIGMOD Record*.

62. Dominich, S. (2008) The Modern Algebra of Information Retrieval. Springer Science & Business Media.

63. (2008) Exploring Network Structure, Dynamics, and Function Using Networkx.

64. Turei, D., Korcsmaros, T. and Saez-Rodriguez, J. (2016) OmniPath: guidelines and gateway for literature-curated signaling pathway resources. Nat Methods, 13, 966–967.

65. Tomlinson, I.P., Novelli, M.R. and Bodmer, W.F. (1996) The mutation rate and cancer. Proc. Natl. Acad. Sci. U. S. A., 93, 14800–14803.

66. Nguyen, B., Fong, C., Luthra, A., Smith, S.A., DiNatale, R.G., Nandakumar, S., Walch, H., Chatila, W.K., Madupuri, R., Kundra, R. et al. (2022) Genomic characterization of metastatic patterns from prospective clinical sequencing of 25,000 patients. Cell, 185, 563–575.e511.

67. Brett, J.O., Spring, L.M., Bardia, A. and Wander, S.A. (2021) ESR1 mutation as an emerging clinical biomarker in metastatic hormone receptor-positive breast cancer. Breast Cancer Res., 23, 85.

68. Fiala, C. and Diamandis, E.P. (2020) Mutations in normal tissues—some diagnostic and clinical implications. BMC Med., 18, 1–9.

69. Laddach, A., Ng, J.C.F. and Fraternali, F. (2021) Pathogenic missense protein variants affect different functional pathways and proteomic features than healthy population variants. PLoS Biol, 19, e3001207.

70. Bianchi, J.J., Zhao, X., Mays, J.C. and Davoli, T. (2020) Not all cancers are created equal: Tissue specificity in cancer genes and pathways. Curr. Opin. Cell Biol., 63, 135–143.

71. Schneider, G., Schmidt-Supprian, M., Rad, R. and Saur, D. (2017) Tissue-specific tumorigenesis: context matters. Nat. Rev. Cancer, 17, 239–253.

72. Yavuz, B.R., Tsai, C.-J., Nussinov, R. and Tuncbag, N. (2023) Pan-cancer clinical impact of latent drivers from double mutations. Communications Biology, 6, 1–14.

73. Avivar-Valderas, A., McEwen, R., Taheri-Ghahfarokhi, A., Carnevalli, L.S., Hardaker, E.L., Maresca, M., Hudson, K., Harrington, E.A. and Cruzalegui, F. (2018) Functional significance of co-occurring mutations in and in breast cancer. Oncotarget, 9, 21444–21458.

74. Luo, Q., Chen, D., Fan, X., Fu, X., Ma, T. and Chen, D. (2020) KRAS and PIK3CA bi-mutations predict a poor prognosis in colorectal cancer patients: A single-site report. Transl. Oncol., 13, 100874.

75. Isnaldi, E., Garuti, A., Cirmena, G., Scabini, S., Rimini, E., Ferrando, L., Lia, M., Murialdo, R., Tixi, L., Carminati, E. et al. (2019) Clinico-pathological associations and concomitant mutations of the RAS/RAF pathway in metastatic colorectal cancer. J. Transl. Med., 17, 137.

76. Yokose, T., Kitago, M., Matsuda, S., Sasaki, Y., Masugi, Y., Nakamura, Y., Shinoda, M., Yagi, H., Abe, Y., Oshima, G. et al. (2020) Combination of KRAS and SMAD4 mutations in formalin-fixed paraffin-embedded tissues as a biomarker for pancreatic cancer. Cancer Sci., 111, 2174–2182.

77. Alon, U. (2007) Network motifs: theory and experimental approaches. Nat. Rev. Genet., 8, 450–461.

78. Iannuccelli, M., Micarelli, E., Surdo, P.L., Palma, A., Perfetto, L., Rozzo, I., Castagnoli, L., Licata, L. and Cesareni, G. (2020) CancerGeneNet: linking driver genes to cancer hallmarks. Nucleic Acids Res., 48, D416–D421.

79. Gorelick, A.N., Sánchez-Rivera, F.J., Cai, Y., Bielski, C.M., Biederstedt, E., Jonsson, P., Richards, A.L., Vasan, N., Penson, A.V., Friedman, N.D. et al. (2020) Phase and context shape the function of composite oncogenic mutations. Nature, 582, 100–103.

80. Hedrick, E., Cheng, Y., Jin, U.-H., Kim, K. and Safe, S. (2016) Specificity protein (Sp) transcription factors Sp1, Sp3 and Sp4 are non-oncogene addiction genes in cancer cells. Oncotarget, 7, 22245-22256.

81. Safe, S., Shrestha, R., Mohankumar, K., Howard, M., Hedrick, E. and Abdelrahim, M. (2021) Transcription factors specificity protein and nuclear receptor 4A1 in pancreatic cancer. World J. Gastroenterol., 27, 6387–6398.

82. Kim, M.P., Li, X., Deng, J., Zhang, Y., Dai, B., Allton, K.L., Hughes, T.G., Siangco, C., Augustine, J.J., Kang, Y.a., et al. (2021) Oncogenic Recruits an Expansive Transcriptional Network through Mutant p53 to Drive Pancreatic Cancer Metastasis. Cancer Discov., 11, 2094–2111.

83. Zhou, Y., Jin, J., Ji, Y., Zhang, J., Fu, N., Chen, M., Wang, J., Qin, K., Jiang, Y., Cheng, D. et al. (2023) TP53 missense mutation reveals gain-of-function properties in small-sized KRAS transformed pancreatic ductal adenocarcinoma. J. Transl. Med., 21, 872.

84. Shi, Y., Hata, A., Lo, R.S., Massagué, J. and Pavletich, N.P. (1997) A structural basis for mutational inactivation of the tumour suppressor Smad4. Nature, 388, 87–93.

85. Subramanian, A., Tamayo, P., Mootha, V.K., Mukherjee, S., Ebert, B.L., Gillette, M.A., Paulovich, A., Pomeroy, S.L., Golub, T.R., Lander, E.S. and Mesirov, J.P. (2005) Gene set enrichment analysis: a knowledge-based approach for interpreting genome-wide expression profiles. Proc. Natl. Acad. Sci. U. S. A., 102, 15545–15550.

86. Mootha, V.K., Lindgren, C.M., Eriksson, K.-F., Subramanian, A., Sihag, S., Lehar, J., Puigserver, P., Carlsson, E., Ridderstråle, M., Laurila, E. et al. (2003) PGC-1alpha- responsive genes involved in oxidative phosphorylation are coordinately downregulated in human diabetes. Nat. Genet., 34, 267–273.

87. Wang, J., Vasaikar, S., Shi, Z., Greer, M. and Zhang, B. (2017) WebGestalt 2017: a more comprehensive, powerful, flexible and interactive gene set enrichment analysis toolkit. Nucleic Acids Res., 45, W130–W137.

88. Singh, A., Nunes, J.J. and Ateeq, B. (2015) Role and therapeutic potential of G-protein coupled receptors in breast cancer progression and metastases. Eur. J. Pharmacol., 763, 178–183.

89. Lappano, R. and Maggiolini, M. (2012) GPCRs and cancer. Acta Pharmacol. Sin., 33, 351–362.

90. Huerta-Reyes, M., Maya-Núñez, G., Pérez-Solis, M.A., López-Muñoz, E., Guillén, N., Olivo-Marin, J.-C. and Aguilar-Rojas, A. (2019) Treatment of Breast Cancer With Gonadotropin-Releasing Hormone Analogs. Front. Oncol., 9, 943.

91. Garrido, M.P., Hernandez, A., Vega, M., Araya, E. and Romero, C. (2023) Conventional and new proposals of GnRH therapy for ovarian, breast, and prostatic cancers. Front. Endocrinol., 14, 1143261.

92. Mishra, P., Banerjee, D. and Ben-Baruch, A. (2011) Chemokines at the crossroads of tumor-fibroblast interactions that promote malignancy. J. Leukoc. Biol., 89, 31–39.

93. Mollica Poeta, V., Massara, M., Capucetti, A. and Bonecchi, R. (2019) Chemokines and Chemokine Receptors: New Targets for Cancer Immunotherapy. Front. Immunol., 10, 379.

94. Lai, X.-R., Wang, C.-L. and Qin, F.-Z. (2022) The mechanism of in lung adenocarcinoma through the SDF-1/CXCR4 pathway. Transl. Cancer Res., 11, 475–487.

95. Liang, K., Liu, Y., Eer, D., Liu, J., Yang, F. and Hu, K. (2018) High CXC Chemokine Ligand 16 (CXCL16) Expression Promotes Proliferation and Metastasis of Lung Cancer via Regulating the NF-κB Pathway. Med. Sci. Monit., 24, 405–411.

96. Liu, H., Yang, Z., Lu, W., Chen, Z., Chen, L., Han, S., Wu, X., Cai, T. and Cai, Y. (2020) Chemokines and chemokine receptors: A new strategy for breast cancer therapy. Cancer Med., 9, 3786–3799.

97. Pajic, M., Herrmann, D., Vennin, C., Conway, J.R., Chin, V.T., Johnsson, A.-K.E., Welch, H.C. and Timpson, P. (2015) The dynamics of Rho GTPase signaling and implications for targeting cancer and the tumor microenvironment. Small GTPases, 6, 123–133.

98. Kimmelman, A.C., Hezel, A.F., Aguirre, A.J., Zheng, H., Paik, J.-H., Ying, H., Chu, G.C., Zhang, J.X., Sahin, E., Yeo, G. et al. (2008) Genomic alterations link Rho family of GTPases to the highly invasive phenotype of pancreas cancer. Proc. Natl. Acad. Sci. U. S. A., 105, 19372–19377.

99. Mohammad, R.M., Li, Y., Muqbil, I., Aboukameel, A., Senapedis, W., Baloglu, E., Landesman, Y., Philip, P.A. and Azmi, A.S. (2019) Targeting Rho GTPase effector p21 activated kinase 4 (PAK4) suppresses p-Bad-microRNA drug resistance axis leading to inhibition of pancreatic ductal adenocarcinoma proliferation. Small GTPases, 10, 367–377.

100. Chen, Z., Wei, X., Dong, S., Han, F., He, R. and Zhou, W. (2022) Challenges and Opportunities Associated With Platelets in Pancreatic Cancer. Front. Oncol., 12, 850485.

101. Mai, S. and Inkielewicz-Stepniak, I. (2021) Pancreatic Cancer and Platelets Crosstalk: A Potential Biomarker and Target. Front Cell Dev Biol, 9, 749689.

102. Zhao, Z.-M., Zhao, B., Bai, Y., Iamarino, A., Gaffney, S.G., Schlessinger, J., Lifton, R.P., Rimm, D.L. and Townsend, J.P. (2016) Early and multiple origins of metastatic lineages within primary tumors. Proc. Natl. Acad. Sci. U. S. A., 113, 2140–2145.

103. Liu, G., Zhan, X., Dong, C. and Liu, L. (2017) Genomics alterations of metastatic and primary tissues across 15 cancer types. Sci. Rep., 7, 13262.

104. Birkbak, N.J. and McGranahan, N. (2020) Cancer Genome Evolutionary Trajectories in Metastasis. Cancer Cell, 37, 8–19.

105. Patel, S.A., Rodrigues, P., Wesolowski, L. and Vanharanta, S. (2021) Genomic control of metastasis. Br. J. Cancer, 124, 3–12.

106. Christensen, D.S., Ahrenfeldt, J., Sokač, M., Kisistók, J., Thomsen, M.K., Maretty, L., McGranahan, N. and Birkbak, N.J. (2022) Treatment Represents a Key Driver of Metastatic Cancer Evolution. Cancer Res., 82, 2918–2927.

107. Theodorou, V., Stark, R., Menon, S. and Carroll, J.S. (2013) GATA3 acts upstream of FOXA1 in mediating ESR1 binding by shaping enhancer accessibility. Genome Res., 23, 12–22.

108. 108. Priestley, P., Baber, J., Lolkema, M.P., Steeghs, N., de Bruijn, E., Shale, C., Duyvesteyn, K., Haidari, S., van Hoeck, A., Onstenk, W., et al. (2019) Pan-cancer whole-genome analyses of metastatic solid tumours. Nature, 575, 210–216.

109. Modrek, A.S., Golub, D., Khan, T., Bready, D., Prado, J., Bowman, C., Deng, J., Zhang, G., Rocha, P.P., Raviram, R. et al. (2017) Low-Grade Astrocytoma Mutations in IDH1, P53, and ATRX Cooperate to Block Differentiation of Human Neural Stem Cells via Repression of SOX2. Cell Rep., 21, 1267-1280.

110. Ohba, S., Kuwahara, K., Yamada, S., Abe, M. and Hirose, Y. (2020) Correlation between IDH, ATRX, and TERT promoter mutations in glioma. Brain Tumor Pathol., 37, 33–40.

111. Saito, Y., Koya, J., Araki, M., Kogure, Y., Shingaki, S., Tabata, M., McClure, M.B., Yoshifuji, K., Matsumoto, S., Isaka, Y. et al. (2020) Landscape and function of multiple mutations within individual oncogenes. Nature, 582, 95–99.

112. Saito, Y., Koya, J. and Kataoka, K. (2021) Multiple mutations within individual oncogenes. Cancer Sci., 112, 483–489.

113. Vasan, N., Razavi, P., Johnson, J.L., Shao, H., Shah, H., Antoine, A., Ladewig, E., Gorelick, A., Lin, T.-Y., Toska, E. et al. (2019) Double mutations in cis increase oncogenicity and sensitivity to PI3Kα inhibitors. Science, 366, 714–723.

114. Iwafuchi-Doi, M., Donahue, G., Kakumanu, A., Watts, J.A., Mahony, S., Pugh, B.F., Lee, D., Kaestner, K.H. and Zaret, K.S. (2016) The Pioneer Transcription Factor FoxA Maintains an Accessible Nucleosome Configuration at Enhancers for Tissue-Specific Gene Activation. Mol. Cell, 62, 79–91.

115. Ciriello, G., Cerami, E., Sander, C. and Schultz, N. (2012) Mutual exclusivity analysis identifies oncogenic network modules. Genome Res., 22, 398–406.

116. Mateo, L., Duran-Frigola, M., Gris-Oliver, A., Palafox, M., Scaltriti, M., Razavi, P., Chandarlapaty, S., Arribas, J., Bellet, M., Serra, V. and Aloy, P. (2020) Personalized cancer therapy prioritization based on driver alteration co-occurrence patterns. Genome Med., 12, 78.

117. Waarts, M.R., Stonestrom, A.J., Park, Y.C. and Levine, R.L. (2022) Targeting mutations in cancer. J Clin Invest, 132.

118. Dunlap, J.B., Leonard, J., Rosenberg, M., Cook, R., Press, R., Fan, G., Raess, P.W., Druker, B.J. and Traer, E. (2019) The combination of NPM1, DNMT3A, and IDH1/2 mutations leads to inferior overall survival in AML. Am. J. Hematol., 94, 913-920.

119. Matsumoto, A., Shimada, Y., Nakano, M., Oyanagi, H., Tajima, Y., Nakano, M., Kameyama, H., Hirose, Y., Ichikawa, H., Nagahashi, M. et al. (2020) RNF43 mutation is associated with aggressive tumor biology along with BRAF V600E mutation in right-sided colorectal cancer. Oncol. Rep., 43, 1853–1862.

120. De, P. and Dey, N. (2019) Mutation-Driven Signals of ARID1A and PI3K Pathways in Ovarian Carcinomas: Alteration Is An Opportunity. Int. J. Mol. Sci., 20.

121. Hopkins, B.D., Goncalves, M.D. and Cantley, L.C. (2020) Insulin-PI3K signalling: an evolutionarily insulated metabolic driver of cancer. Nat. Rev. Endocrinol., 16, 276–283.

122. Tanaka, H., Takizawa, Y., Takaku, M., Kato, D., Kumagawa, Y., Grimm, S.A., Wade, P.A. and Kurumizaka, H. (2020) Interaction of the pioneer transcription factor GATA3 with nucleosomes. Nat. Commun., 11, 4136.

123. Zhou, Y., Zhao, J., Fang, J., Martin, W., Li, L., Nussinov, R., Chan, T.A., Eng, C. and Cheng, F. (2021) My personal mutanome: a computational genomic medicine platform for searching network perturbing alleles linking genotype to phenotype. Genome Biol., 22, 53.

124. Liu, C., Han, Z., Zhang, Z.-K., Nussinov, R. and Cheng, F. (2021) A network-based deep learning methodology for stratification of tumor mutations. Bioinformatics, 37, 82–88.

125. Chang, L., Jung, N.Y., Atari, A., Rodriguez, D.J., Kesar, D., Song, T.Y., Rees, M.G., Ronan, M., Li, R., Ruiz, P. et al. (2023) Systematic profiling of conditional pathway activation identifies context-dependent synthetic lethalities. Nat Genet, 55, 1709–1720.

