## Supplementary File for "Co-occurring Mutations in Different Genes Can Fuel Oncogenic Signaling and Serve as Metastatic Tumor Markers"

**Supplementary Material**

**FP Growth Tree Construction**

*Built*

Workflow for analyzing mutation patterns in a dataset, specifically BreastMetastatic cohort, using frequent pattern mining, which is a common task in bioinformatics and data mining. This process appears to use the **mlextend** library's **FreqItems** and **AssociationRules** functions, which are likely Python implementations for mining frequent itemsets and association rules, respectively. Here's a more detailed explanation based on the FP-Tree (Frequent Pattern Tree) algorithm and the **mlextend** library:

*Input Data*

The workflow begins with a binary alteration matrix from the BreastMetastatic cohort. This matrix likely has cases as rows and mutations as columns, as binary format indicates the lacking/having of mutations corresponds with 0/1.

*Frequent Itemset Mining*

The **mlextend.FreqItems** function processes the binary matrix to find frequent mutation patterns. The function is parameterized by **minSupport** set to **0.001**, which is the minimum frequency a pattern must have to be considered **frequent**.

This process identifies sets of mutations that commonly occur together in the dataset. The frequent itemsets are parsed from the NULL tree, which is a reference to the initial state of the FP-Tree where no patterns have yet been identified.

*Association Rule Mining*

The frequent itemsets obtained are used to generate association rules with the **AssociationRules** function from **mlextend**.

This function looks for mutations that are highly associated with each other, based on a **minTreshold** parameter. This threshold determines how strong the association needs to be for a rule to be considered. It's noted that unlike a p-value, the confidence of these rules is inversely related to the p-value from statistical tests.

*Tree Construction*

Based on the association rules, new branches (paths) are generated and added to the NULL (ROOT) node of the FP-Tree. Only branches representing associated pairs are added.

This step constructs a tree where the paths represent the common mutations between different cases, revealing how certain mutations (ancestors) are associated with other mutations (descendants) within the dataset.

*Tree Visualization*

The layout and theme of the tree are adjusted according to the characteristics of the pairs/branches:

- The **EdgeName** is labeled with the **PairCounts**, which likely indicates the number of times that pair of mutations has been found associated.
- The **NodeSize** represents the frequency of the mutation or alteration, which could be visualized by larger nodes indicating more frequent mutations.
- The **EdgeWidth** is determined by the **PairCounts**, where a wider edge indicates a more frequently occurring pair of mutations.
- The **EdgeStyle** (dashed or linear) differentiates between mutation pairs (AlterationPair) and edges that stem directly from the root of the tree.
- The **ColorTones** used in the visualization distinguish between ancestors (*ANCE*, dark tones) and descendants (*CONS*, light tones) to show the directionality of associations.
- The GitHub link provided -> https://github.com/ugur0sahin/FMPSeeker

contain the implementation of this workflow or additional documentation on how the FP-Tree algorithm is adapted for this specific use case.

FP-Tree algorithm to find frequent patterns and strong association rules in a binary matrix representing mutation data from breast cancer metastatic cases. The final goal is to construct a tree visualization that meaningfully represents the relationships between different mutations, providing insights into the underlying patterns of mutation co-occurrence in the data.

*Workflow:*

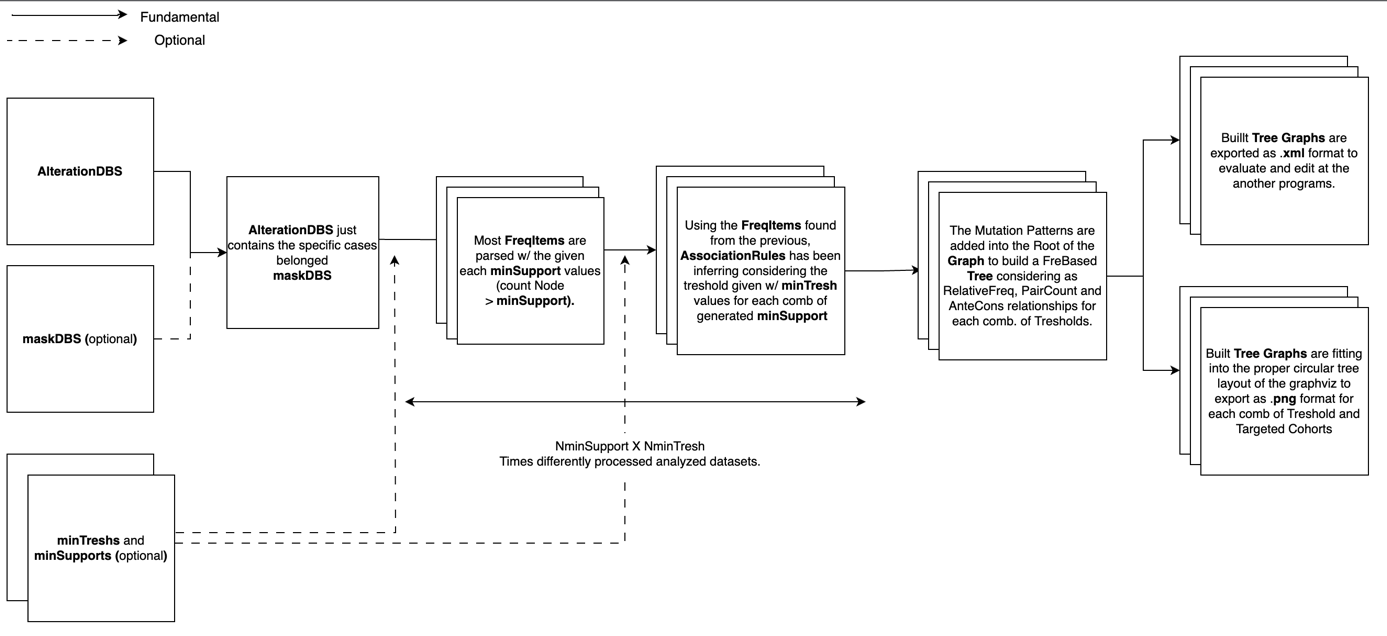

**Supplementary Figures**

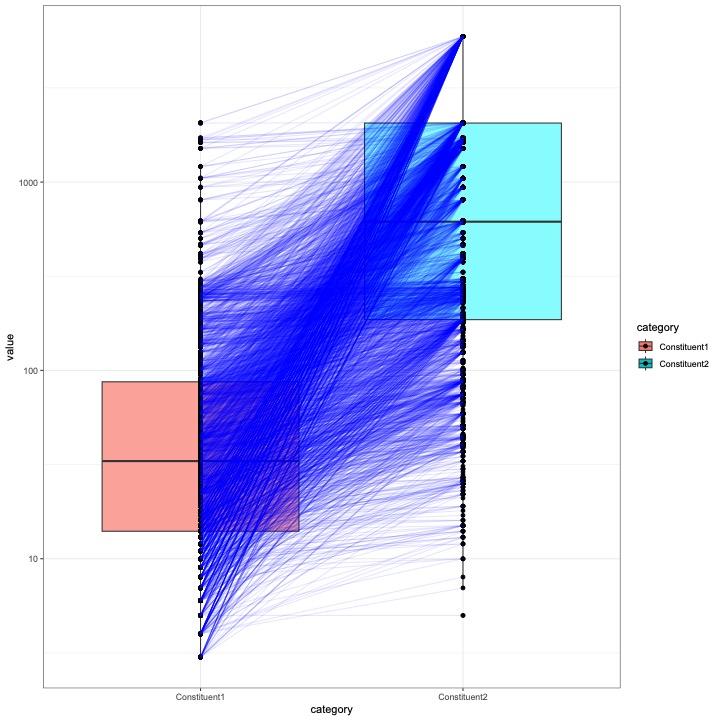

**Figure S1:** Box plot showing the frequency distribution of double mutation constituents with connected dots. The lines depicts indicates the co-occurrence of a frequent mutation with a relatively rare one.

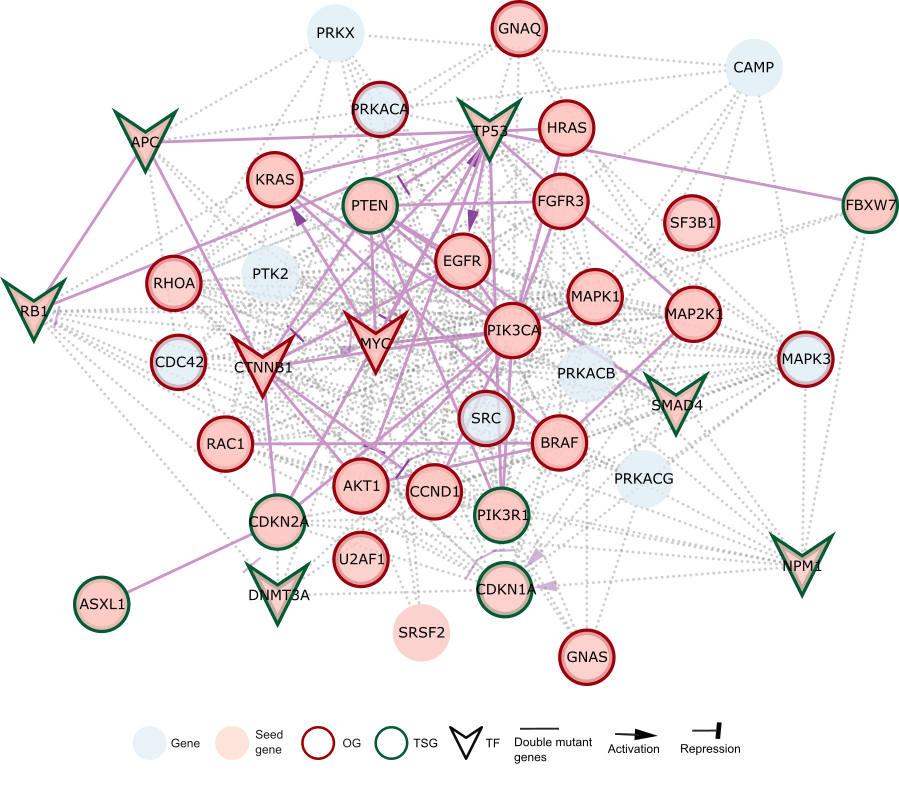

**Figure S2:** Pancreas specific subnetwork. There are 40 nodes 267 edges in the subnetwork obtained from Omni Path PPI by Page Rank Algorithm with 23 seed genes.

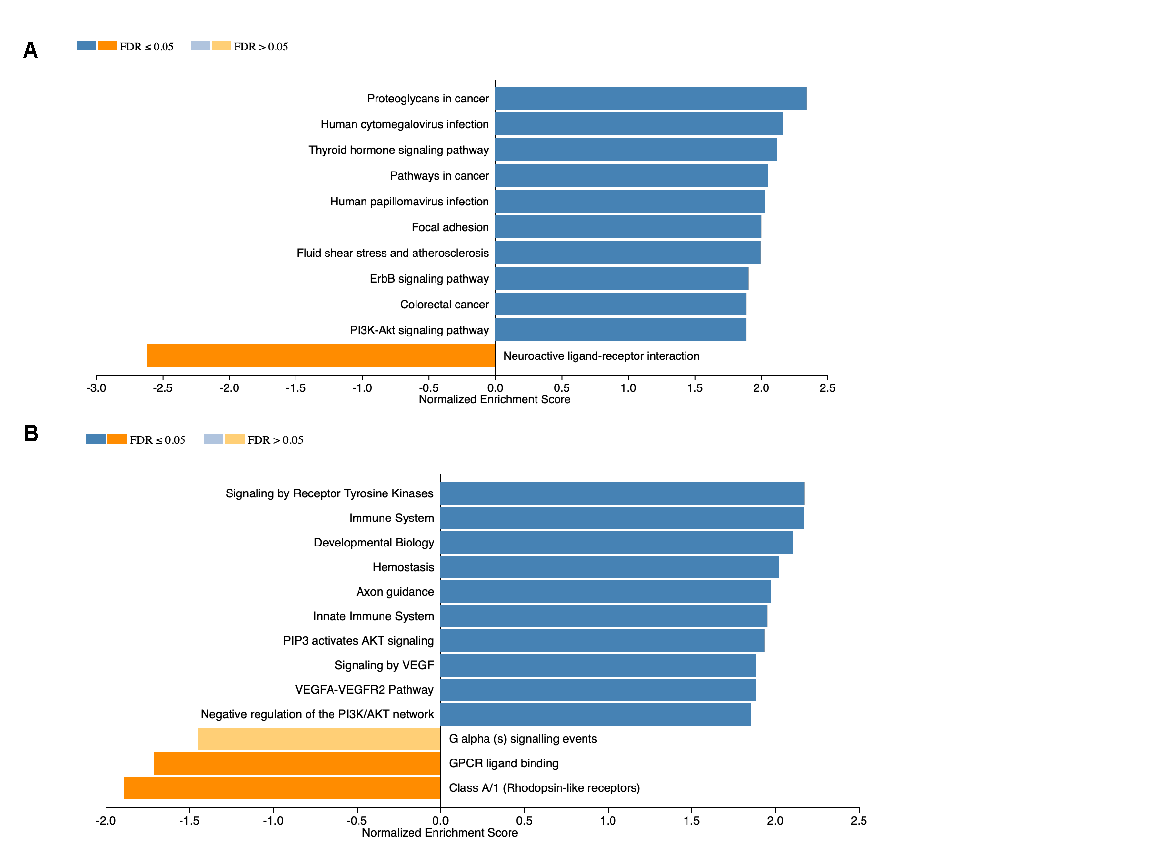

**Figure S3:** Gene set enrichment analysis of breast cancer specific subnetwork in (A) KEGG , (B) Reactome.

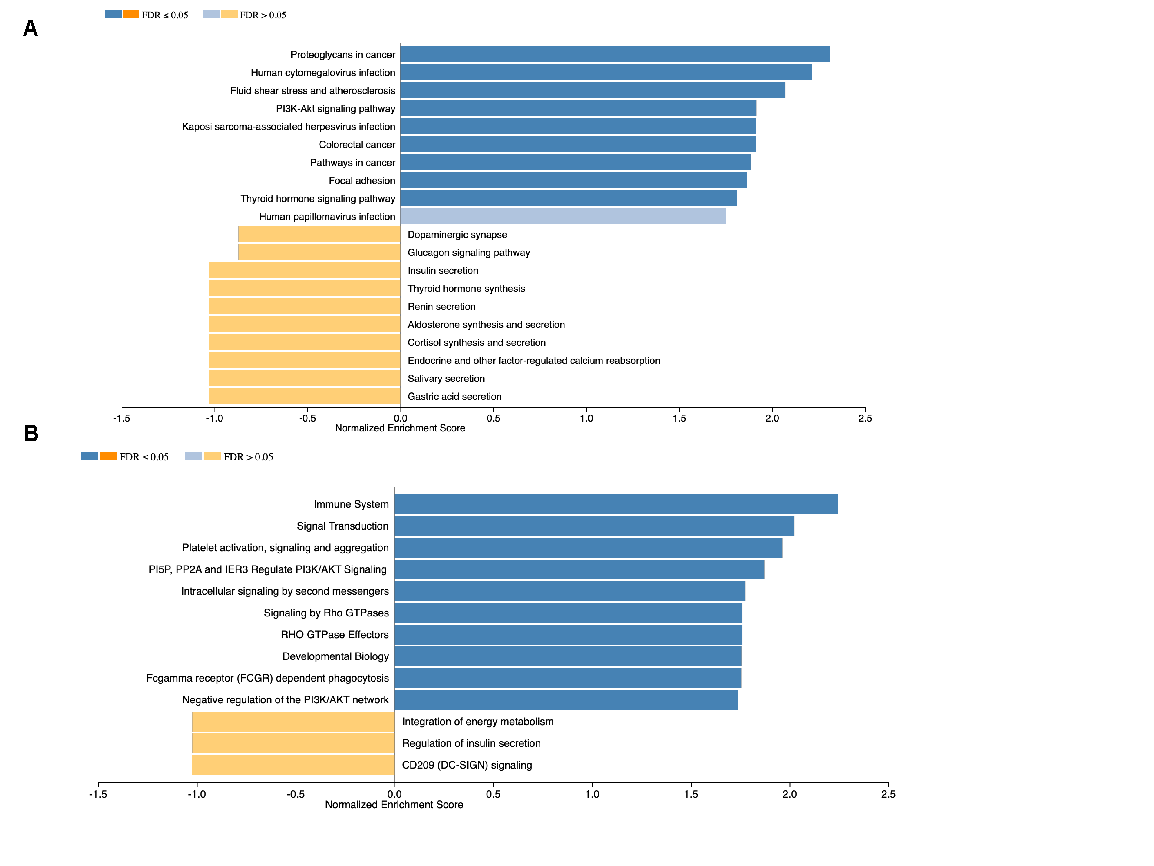

**Figure S4:** Gene set enrichment analysis of pancreas cancer specific subnetwork in (A) KEGG, (B) Reactome.

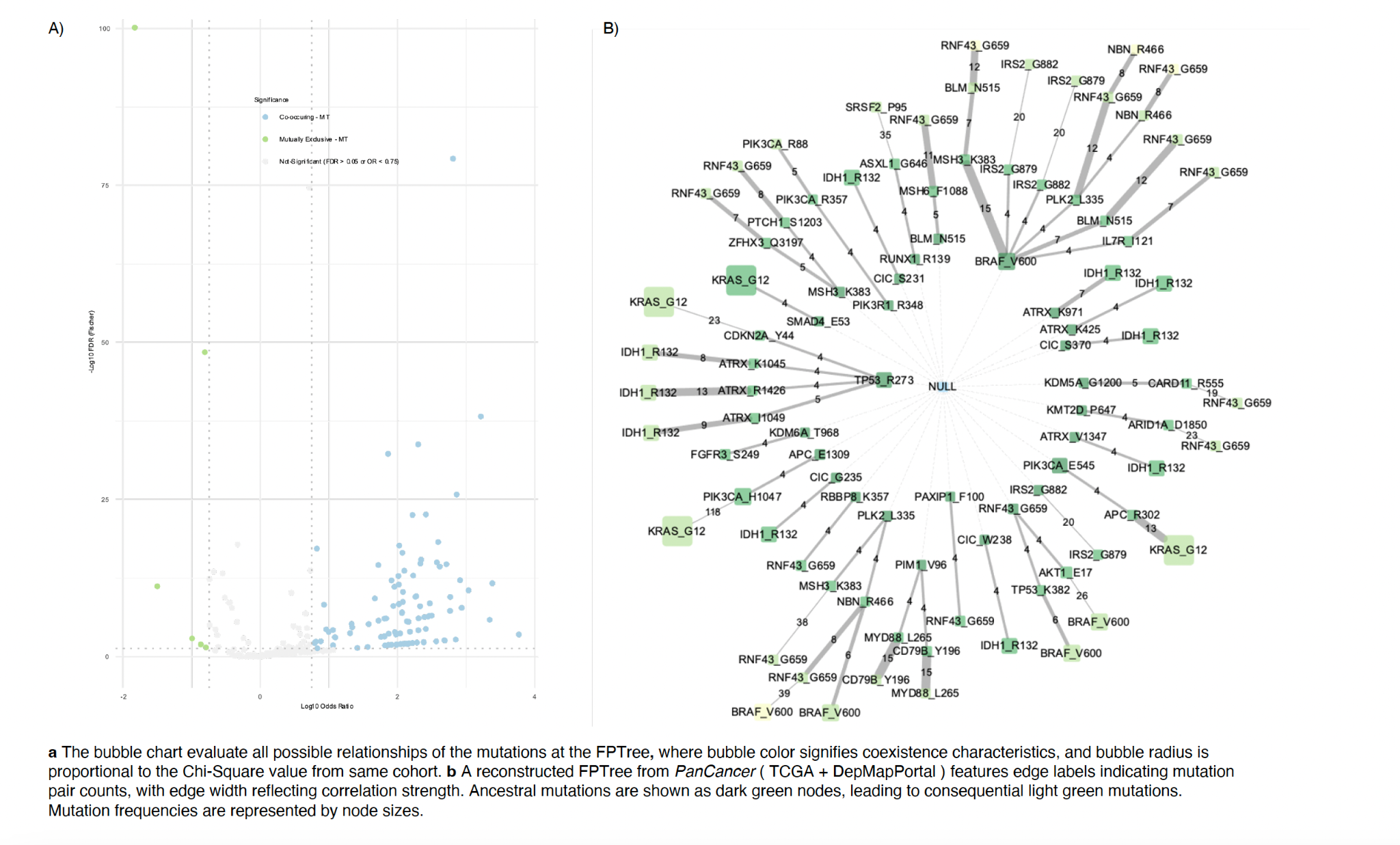

**Figure S5: (A**)The Volcano plot of pairs of same cohort, log10 Odds Ratio against -log10 Fisher’s p-value, with non-significant pairs (σ > 0.05, |Odds Ratio| > 0.75) in gray. (B) reconstructed FPTree from pan-cancer metastatic cohort features edge labels indicating mutation pair counts, with edge width reflecting correlation strength. Ancestral mutations are shown as dark green nodes, leading to consequential light green mutations. Mutation frequencies are represented by node sizes.
